# A tripartite cytolytic toxin formed by *Vibrio cholerae* proteins with flagellum-facilitated secretion

**DOI:** 10.1101/2021.06.20.449157

**Authors:** Aftab Nadeem, Raghavendra Nagampalli, Eric Toh, Athar Alam, Si Lhyum Myint, Thomas V. Heidler, Mitesh Dongre, Nikola Zlatkov, Hudson Pace, Fouzia Bano, Anders Sjöstedt, Marta Bally, Bernt Eric Uhlin, Sun Nyunt Wai, Karina Persson

## Abstract

The protein MakA was discovered as a motility-associated secreted toxin from *Vibrio cholerae*, Here, we show that MakA is part of a gene cluster encoding four additional proteins: MakB, MakC, MakD and MakE. The MakA, MakB and MakE proteins were readily detected in culture supernatants of wild type *V. cholerae* whereas secretion was very much reduced from a flagellum deficient mutant. Crystal structures of MakA, MakB and MakE revealed structural relationship to a superfamily of bacterial pore-forming proteins. Cloning and expression of MakA/B/E in *Escherichia coli* resulted in toxicity of the bacteria towards *Caenorhabditis elegans* used as a predatory organism model. None of these Mak proteins alone or in pairwise combinations were cytolytic but an equimolar mixture of MakA, MakB and MakE acted as a tripartite cytolytic toxin in vitro causing lysis of erythrocytes and cytotoxicity on cultured human colon carcinoma cells. Formation of oligomeric complexes on liposomes was observed by electron microscopy. Oligomer interaction with membranes was initiated by MakA membrane binding followed by MakB and MakE joining in formation of a pore structure. A predicted membrane insertion domain of MakA was shown by site-directed mutagenesis to be essential for toxicity towards *C. elegans*. Bioinformatic analyses revealed that the *makCDBAE* gene cluster is present as a novel genomic island in the vast majority of sequenced genomes of *V. cholerae* and the fish pathogen *V. anguillarum*. We suggest that the hitherto unrecognized cytolytic MakA/B/E toxin can contribute to *Vibrionaceae* fitness and virulence potential in different host environments and organisms.

**Significance Statement:** *Vibrio cholerae*, responsible for outbreaks and pandemics of cholera disease, is a highly motile organism by virtue of a single flagellum. We describe that the flagellum facilitates the secretion of three *V. cholerae* proteins encoded by a hitherto unrecognized novel genomic island. The proteins MakA/B/E can form a tripartite cytolytic toxin that lyses erythrocytes and is cytotoxic to cultured human cells. A structural basis for the Mak protein cytolytic activity was obtained by X-ray crystallography. Flagellum-facilitated secretion, remarkably ensuring spatially co-ordinated delivery of Mak proteins, revealed a new role for the *V. cholerae* flagellum considered of particular significance for the bacterial environmental persistence. Our findings will pave the way for the development of new diagnostics and therapeutic strategies against pathogenic *Vibrionaceae*.

## INTRODUCTION

Vibrios (*Vibrionaceae* strains) are comma- or rod-shaped, motile Gammaproteobacteria found in aquatic habitats and in association with eukaryotes. *Vibrio cholerae* is known as the cause of cholera, an infectious disease causing watery diarrhea that can lead to fatal dehydration (1). Cholera outbreaks are frequent in low and middle-income countries, and the major sources are contaminated water and poor sanitation, often in combination with natural disasters. The disease is caused by a few serogroups of *V. cholera*e, and the main factor behind the cholera symptoms is the cholera toxin (CT), an A-B toxin encoded by genes located on a prophage mobile genetic element (CTXφ) that induce severe disruption of intestinal cell function, leading to the watery, secretory diarrhea (2, 3). Most serogroups do not cause cholera, as they do not possess the genes for CT, but they cause other diseases, e.g., skin, wound, and gastrointestinal infections as well as bacteremia and septicemia (4-6). The natural reservoirs of *V. cholerae* are aquatic sources such as rivers, brackish waters, and estuaries and are often associated with copepods or other zooplankton, aquatic plants and shellfish (7). It is of great interest to elucidate the actual factors and mechanisms enabling *V. cholerae* and other *Vibrionaceae* to survive and persist in natural environments with challenging growth conditions and predators (8).

Like many other Gram-negative bacteria, *V. cholerae* is motile by virtue of flagella, the rotating motor organelles that enable bacteria to move. While some bacteria have multiple flagella, *V. cholerae* possesses a single polar flagellum. The flagellum export machinery and the virulence-associated type-III secretion system (denoted as fT3SS and vT3SS, respectively) are suggested to share a common ancestor (9), explaining their similar structure and molecular structure organization. The vT3SS allows the delivery of effector proteins through a hollow channel directly into the eukaryotic host cell (10). The fT3SS has a similar structure, including a channel through which flagellar proteins are transported during flagellum assembly. In the bacterial cytoplasm, effectors secreted by the vT3SS are stabilized by chaperones to prevent aggregation. These chaperones are often encoded by genes adjacent to those encoding the effectors (11). Flagellar proteins are similarly protected by chaperones before they are transported to the growing distal end of the flagellum (12).

We earlier established *Caenorhabditis elegans* as a predatory organism model for identifying and assessing factors other than CT from *V. cholerae* that may contribute to bacterial survival and persistence in the environment (13). With this model, we discovered a novel cytotoxin, MakA (<u>m</u>otility-<u>a</u>ssociated <u>k</u>illing factor <u>A</u>). The *makA* gene locus was not previously known to be involved in the virulence of *V. cholerae*, but we demonstrated that MakA is an essential factor for the cytotoxic activity of *V. cholerae* in both *C. elegans* and *Danio rerio* (zebrafish) (14). We further demonstrated that secretion of MakA occurs via the flagellum channel in a manner that is novel in *V. cholerae*.

Our crystal structure of MakA revealed structural similarities to ClyA (14), the pore-forming protein first identified in non-pathogenic *Escherichia coli* (15, 16) and subsequently also found in *Salmonella enterica* Serovars Typhi and Paratyphi A (17). ClyA from *E. coli* is expressed from a monocistronic operon and functions as a toxin on its own upon release via membrane vesicles (16, 18, 19). In contrast, *V. cholerae* MakA is expressed from an operon that encodes four additional proteins for which the roles are not known. MakA is structurally related also to two proteins from the Gram-positive bacterium *Bacillus cereus*, the hemolysin BL binding component B (HBL-B) and the NheA component of the Nhe non-hemolytic enterotoxin. Both of these are considered components of tripartite toxins (20). Recently, a tripartite toxin, the AhlABC toxin, was identified and structurally characterized as a pore-forming toxin in the Gram-negative species *Aeromonas hydrophila*, and it is important to note that the structure of soluble AhlB shares the same general structure as earlier described for MakA (21). In addition, a similar toxin complex of three proteins, SmhABC from *Serratia marcescens*, was recently reported (22). However, if and how the Ahl and Smh proteins are released during normal growth, or if there is a dedicated secretion system, remain unclear.

Here we identify the proteins from the five *V. cholera* genes, *vca0880-vca0884*, that are co-expressed from the operon denoted *makDCABE*. The structure and function of three of the proteins, MakA, MakB and MakE were characterized in detail. Crystal structure analyses revealed that the MakA, MakB and MakE proteins all share structural features with a group of bacterial pore-forming toxins (PFTs) with defined membrane-penetrating domains. Our *in vitro* studies revealed that an equimolar combination of the MakA/B/E proteins acted as a tripartite cytotoxin causing efficient lysis of red blood cells and cytotoxicity to human cells. Furthermore, examination of a large number of bacterial genome sequences revealed that the *mak* gene cluster is present in many *V. cholerae* and other *Vibrionaceae* strains. These include *Vibrio anguillarum*, an inhabitant of estuarine and marine coastal ecosystems worldwide and the etiological agent of classical vibriosis in warm- and cold-water fish species (23). The identification and structural characterization of the Mak proteins in *V. cholerae* presented here reveals a hitherto unrecognised potential of many pathogenic *Vibrionaceae* strains to produce the tripartite Mak cytolytic toxin.

## RESULTS

### Expression and secretion of MakA, MakB and MakE proteins from *Vibrio cholerae*

MakA is encoded from a gene cluster in *V. cholerae* O1 strain A1552 where the three genes located upstream of *makA* were denoted *makD, makC* and *makB* (14). An additional gene, here denoted *makE*, is located downstream of *makA*. The *mak* gene cluster thereby includes five genes in an operon transcribed in the direction *makD –> makC* –> *makB –> makA –> makE* (**Fig. 1A**). By molecular cloning and mutagenesis of each gene, we analyzed the nature of the predicted proteins. Tests with *V. cholerae* mutants defective in *makA* or *makB* demonstrated a clear attenuation of toxicity in *C. elegans* whereas the effect of Δ*makD* or Δ*makC* mutations was minimal (14).

**Figure 1.**
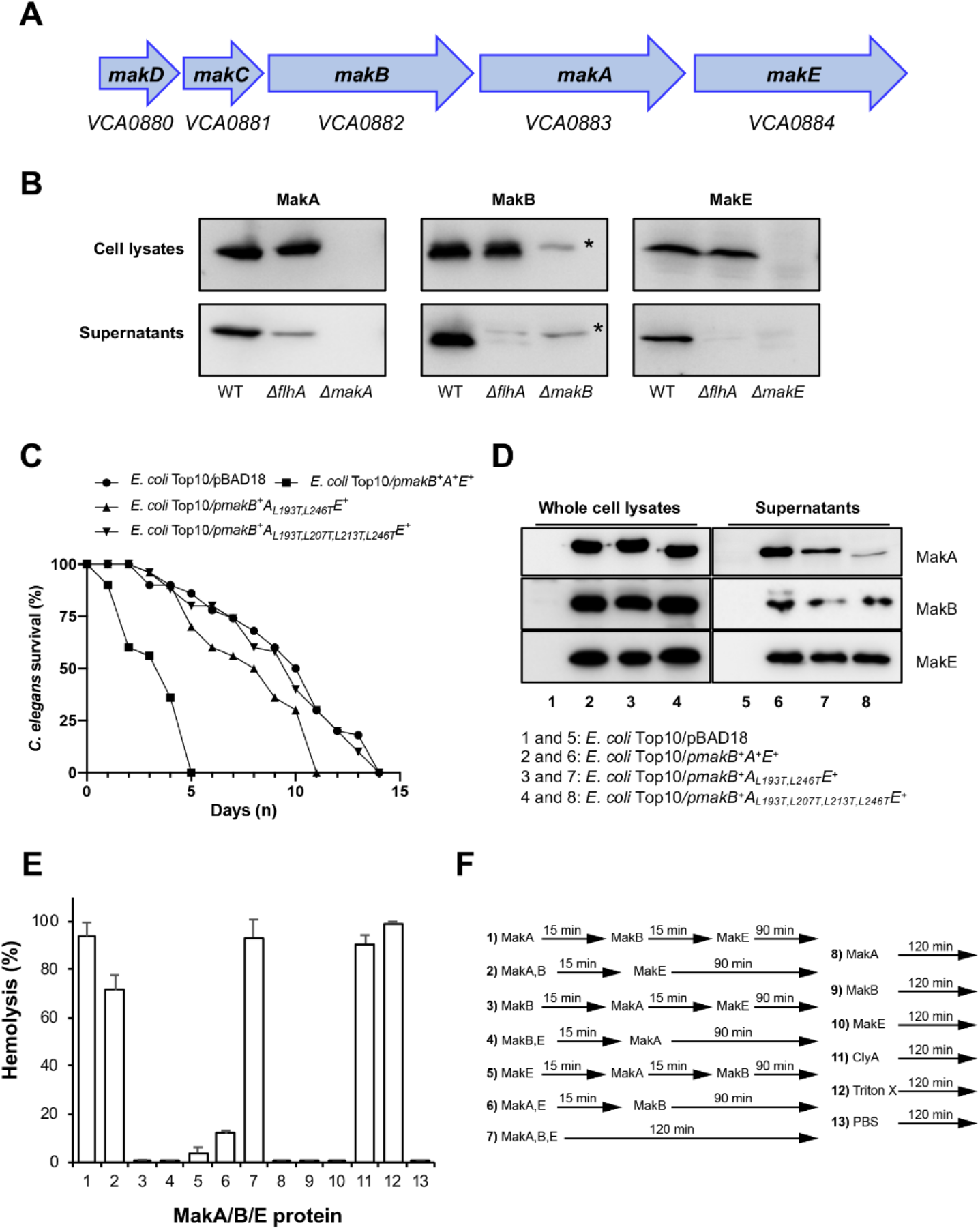
*Vibrio cholerae* secretion of Mak proteins forming a tripartite cytolytic toxin. ***(A)*** Gene organization of the *mak* operon in the genome of *Vibrio cholerae* O1 El Tor strains A1552 and N16961. The gene loci *vca0880, vca0881, vca0882* and *vca0883* were earlier denoted *makD, makC, makB* and *makA*, respectively(14). Here we identify *vca0884* as an additional *mak* gene, *makE*. ***(B)*** Flagella-dependent secretion of Mak proteins. Western immunoblot detection of MakA, MakB, and MakE in whole cell lysates and in the supernatant. The study was done on wild-type *V. cholerae* A1552, Δ*flhA*, Δ*makA, ΔmakB, and ΔmakE* mutants. The proteins were detected with antisera raised against MakA, MakB and MakE respectively. ***(C)*** *C. elegans* survival upon feeding on *E. coli* Top10 strains harboring different *mak* genes. Data represent average survival percentage of three separate experiments. ***(D)*** Western blot analysis of MakA, MakB, or MakE protein secreted in the culture supernatant or from bacterial whole cell lysate of the *E. coli* Top10 strains overexpressing native or MakA head mutant protein together with MakB and MakE. ***(E-F)*** Lysis of erythrocytes by Mak proteins in solution. 250 nM of each of MakA, MakB and MakE were added individually in sequential order or in combination to a solution of human erythrocytes (2% of whole blood) and incubated for 120 min at 37 °C. The cytolytic *E. coli* protein ClyA (250 nM) was used as a positive control, and PBS was the negative control. After centrifugation, the supernatants were monitored spectrophotometrically for released haemoglobin, by measurement of absorbance at 545 nm, as an indicator of red blood cell lysis. Data are from three independent experiments.

We have demonstrated that MakA, which is comprised of 369 amino acids, is mainly secreted via the *V. cholerae* flagellum (14). To test if MakB and MakE (354 and 353 amino acids, respectively) also are secreted, we performed immunoblot analyses of samples from cell lysates and supernatants using anti-MakA, anti-MakB, and anti-MakE antisera (**Fig. 1B**). Similar to the earlier findings with MakA, both MakB and MakE were detected in the supernatant. Furthermore, when we monitored MakA, MakB and MakE expressed by a flagellum-deficient (Δ*flhA*) mutant of *V. cholerae* strain A1552, the secretion of all three proteins was drastically reduced (**Fig. 1B**). We conclude that secretion of these three Mak proteins is facilitated by the *V. cholerae* flagellum as earlier demonstrated for MakA (14). It should be noted that MakA secretion occurred to a level of about 10% even from the Δ*flhA* mutant derivative(14).

We also monitored the expression and secretion of Mak proteins in *V. cholerae* A1552 derivatives mutated in each of the *mak* genes (**Supplementary Fig. 1**). Immunodetection of the cytoplasmic protein CRP was used as a reference and a non-secreted control. There was a strong reduction in the cellular level of MakA in the *ΔmakB* mutant, and consequently, very little MakA was secreted. Also, the *ΔmakC, ΔmakD*, and *ΔmakE* mutants showed a somewhat lower cellular level of MakA, and accordingly, the levels were lower in the supernatants (**Supplementary Fig. 1A**, lanes 3, 9-12). The low amount of MakA in the *ΔmakB* mutant remained even after the strain was complemented *in trans* with a plasmid expressing MakB (**Supplementary Fig. 1E**). Similarly, MakB expression and secretion levels were much reduced in the *ΔmakC* mutant (**Supplementary Fig. 1B & D**, lanes 4 & 10). In addition, the amount of MakB in the Δ*makC* mutant remained low in the strain complemented *in trans* with a plasmid expressing MakC (**Supplementary Fig. 1F**). We interpreted the results as an indication that deletions of *makB* or *makC* caused strong polarity effects, in particular on the expression of their immediate downstream neighboring genes.

We did not detect any secretion of the MakC protein (**Supplementary Fig. 1G**) and we consider that both MakC and MakD have accessory roles and remain in the bacterial cytoplasm or periplasm.

The Vibrio flagellum is covered by a sheath, an extension of the outer membrane that contains LPS and surrounds the filament (24, 25). Secretion of the anti-sigma factor FlgM through the filament was demonstrated, suggesting that there may be a sheath opening at the flagellar tip. Recent studies suggest that the H-ring is essential for outer membrane penetration and assembly of the polar flagellum in Vibrio spp. (26). Vibrios lacking FlgT (a component of the H-ring) were shown to synthesize some flagella that failed to penetrate the outer membrane, forming periplasmic flagella, and it was considered that the H-ring might play a role in sheath formation. In the absence of any known mutant lacking the sheath *per se*, we decided to test if secretion and expression of the MakA/B/E proteins would be affected by a Δ*flgT* mutation in *V. cholerae* strain A1552. The results showed that the Δ*flgT* mutation did not cause any difference in secretion and expression of the MakA/B/E proteins compared to the wild type strain (**Supplementary Fig. 2**). These results suggest that the proposed role of the H-ring to facilitate the outer membrane penetration of the polar sheathed flagellum is not essential for the secretion of the Mak proteins.

The finding that the three proteins MakA, MakB and MakE are secreted in a similar flagella-facilitated manner prompted us to test if they can also be secreted from *E. coli* as was shown earlier with MakA (14). A plasmid clone with an inducible transcriptional promoter that encoded the *makB*^*+*^ *makA*^*+*^ and *makE*^*+*^ genes from the *mak* operon was obtained. We tested if the construct would mediate a toxic effect on *C. elegans*, similar to what we found when MakA was recognized as a toxin in this model host organism for *V. cholera*e (14). As shown in **Fig. 1C** and **Fig. 1D** (lanes 2), the MakA/B/E expression in *E. coli* caused killing of *C. elegans*, and all three proteins were readily detected as secreted in the supernatant. We can thereby also conclude that the presence of the other two proteins, MakC and MakD, from the *mak* operon is not essential for the secretion of MakA/B/E and their toxin activity.

### The MakA/B/E tripartite protein combination has hemolytic and cytotoxic activity

We purified recombinant MakA, MakB and MakE for further characterization. To test if any of the individual Mak proteins or bipartite/tripartite combinations thereof would be cytolytic, we put samples onto blood agar plates and monitored for erythrocyte lysis (**Supplementary Fig. 3A)**. As a positive control, we used the earlier characterized cytolytic protein ClyA from *E. coli* (15, 16). We observed a clear hemolytic effect of the equimolar combination of the tripartite combination (MakA/B/E), but none for the individual proteins or bipartite combinations.

The cytolytic activity of the MakA/B/E tripartite was further assessed by quantitative hemolysis assays with erythrocytes in solution and with a 120 min treatment at 37°C (**Fig. 1E-F**). To determine if adding the individual MakA/B/E proteins to the tripartite combination would be of importance and affect the activity in the hemolysis assay, the proteins (each at 250nM) were introduced in a series of different sequential orders during the 120 min treatment **(Fig. 1E-F)**. Maximum hemolytic activity was observed with the order MakA -> MakB -> MakE with intervals of 15 min (**Fig. 1E, column 1**) and when all three components of the tripartite were mixed prior to their addition to the erythrocytes (**Fig. 1E, column 7**). Notably, the hemolytic activity of the equimolar MakA/B/E tripartite was similar to that observed with 250 nM of the ClyA protein (**Fig. 1E, column 11**). A clearly detected hemolysis, albeit to about 10-15% lower level, was also observed when MakA and MakB were first introduced as a premixed sample before MakE was added as the last component (**Fig. 1E, column 2**). Interestingly, when MakB was added first, followed by MakA and finally MakE, there was no hemolysis detected within the 120 min assay (**Fig. 1E, column 3**). Similarly, when MakB and MakE were first introduced as a premixed sample before MakA was added 15 min later, there was no detectable hemolysis (**Fig. 1E, column 4**). A low level of hemolysis (5-10%) was detected when the proteins were added separately in the order MakE->MakA->MakB or in the order MakAE->MakB (**Fig. 1, columns 5 and 6**, respectively). Finally, neither of the three Mak proteins tested alone caused any detectable hemolysis in this assay (**Fig. 1, columns 8-10**). We conclude from these results that all three proteins MakA, MakB and MakE were required for the cytolytic activity and that there was a clear dependence of in which order the proteins were introduced. Only when MakA was added first alone, or in a premix with MakB, did the tripartite result in high level hemolysis.

In addition to displaying hemolytic activity, the MakA/B/E tripartite could be visualized on the red blood cell surfaces, as recorded by confocal microscopy (**Fig. 2A**). The binding kinetics of the Mak tripartite to a lipid membrane in real time was studied using supported synthetic lipid membrane (SLM) bilayers of complex composition and analysis in a Quartz Crystal Microbalance with Dissipation monitoring (QCM-D) of the protein adsorption (**Supplementary Fig. 3B**), confirming protein binding to the bilayer surface. The proteins were stably bound and could not be washed off upon rinsing. Liposomes prepared from an *E. coli* total lipid extract and analysis by transmission electron microscopy were used to investigate if the MakA/B/E tripartite complex or its individual components would form recognizable oligomeric or pore-like assemblies in the presence of lipid membranes. Formation of oligomeric assemblies were indeed observed with the MakA/B/E tripartite complex (**Fig. 2B**). Among the individual Mak proteins, only MakA formed seemingly well-organized star-shaped oligomeric structures.

**Figure 2.**
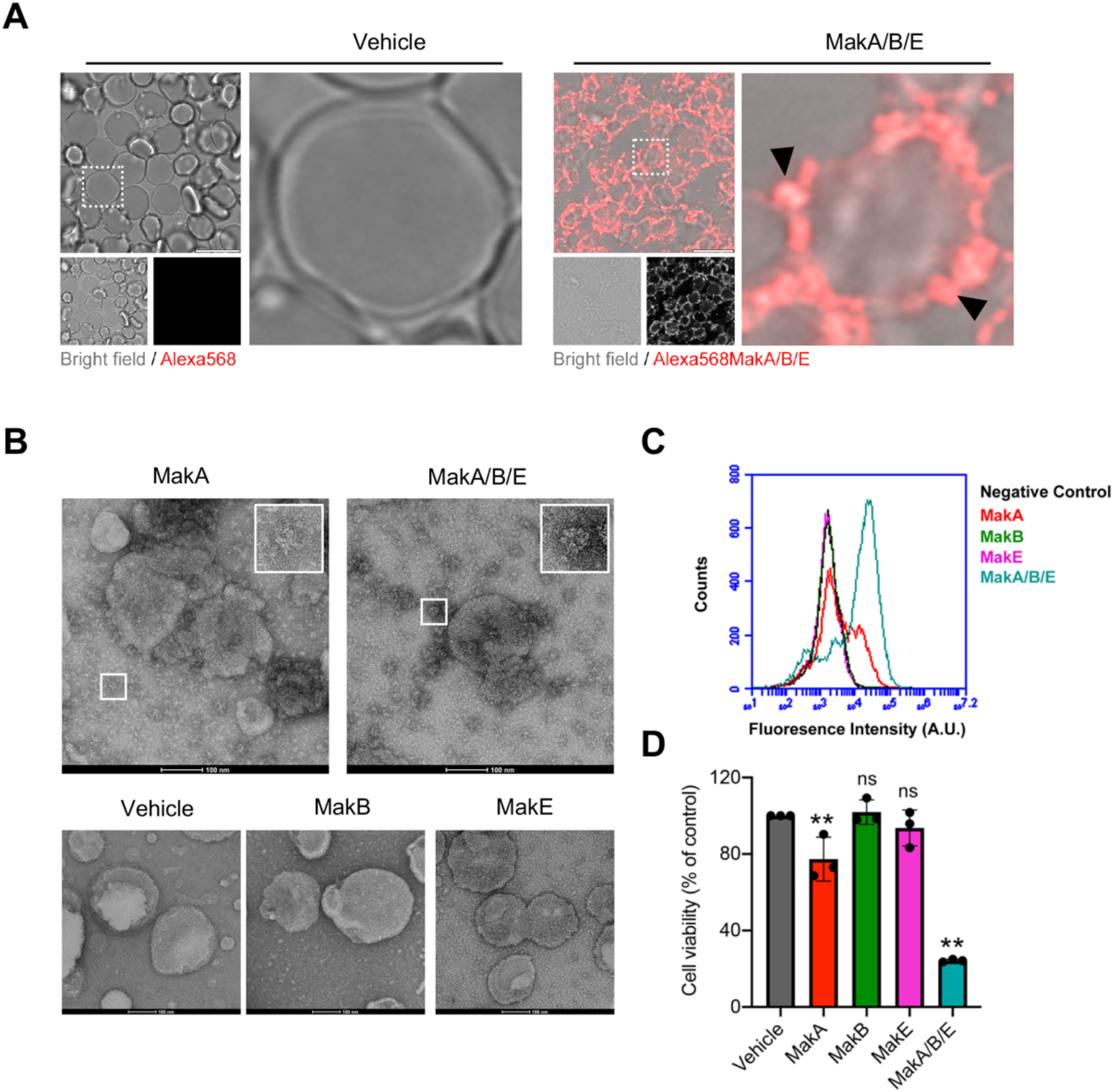
Activity of the tripartite MakA, MakB and MakE cytotoxin on human cells. ***(A)*** Human red blood cells (1% from whole blood) in phosphate buffer saline were treated with the Alexa568 labelled MakA, MakB and MakE tripartite protein combination (125 nM of each). Binding and membrane accumulation of the Alexa568 labelled proteins were assessed by confocal laser scanning microscopy. Arrowheads (black) indicate membrane accumulation of Alexa568-MakA/B/E tripartite complex. Scale bars represent 10 µm. ***(B)*** *E. coli* total cell lipid extract liposomes were treated with vehicle (Tris 20mM), MakA, MakB, MakE or MakA/B/E tripartite complex for 90 min and stained with 1.5% uranyl acetate solution. Micrographs were captured with transmission electron microscopy (TEM). Inset indicates star-shaped oligomeric structure for MakA and larger complexes for the MakA/B/E tripartite complex. Scale bars represent 100 nm. ***(C)*** Human colon cancer cell line, CaCO2 cells were treated with Alexa-568 labelled preparations (at 250 nM) of MakA, MakB and MakE both individually and in the tripartite combination at equimolar concentrations. The cell-associated Alexa-568 labelled protein was assessed after 24 h treatment by flow cytometry analysis. ***(D)*** Effect of Mak proteins on the viability of colon cancer cells. CaCO2 cells were treated with the MakA, MakB and MakE proteins (at 250 nM), individually and in the tripartite combination, for 24 h and toxicity was assessed by the MTS cell viability assay. Data points represent four biologically independent experiments; bar graphs show mean ± s.d. Significance was determined from biological replicates using one-way analysis of variance (ANOVA) with Dunnett’s multiple comparisons test. *p<0.05, **p<0.01, ns = not significant.

To investigate if the individual proteins and/or the MakA/B/E tripartite would bind to mammalian epithelial cells, we used a cell line of human colon cancer cells (CaCO2 cells) and tested for interaction with Alexa568-labelled preparations of the MakA, MakB, and MakE proteins. Flow cytometry analysis was used to monitor the label representing binding and/or uptake of the Alexa568-labeled Mak proteins. The results indicated that a distinctly higher amount of cells were labelled with the MakA/B/E tripartite combination than with any of the three proteins tested separately (**Fig. 2C**). Of the three Mak proteins individually tested, MakA showed higher binding/uptake than MakB or MakE. We also assessed the viability of the CaCO2 cells upon 24 h treatment with the Mak proteins. The results from the cell viability assay showed that the MakA/B/E tripartite combination displayed the most pronounced cytotoxic activity, resulting in a loss of viability in about 80% of the CaCO2 cells (**Fig. 2D**). A lower but still significant degree of cytotoxicity was seen with MakA alone.

To determine if the MakA/B/E tripartite disrupted the epithelial cell intracellular structures, CaCO2 cells treated with the tripartite combination were stained for: i) actin filaments using Phalloidin-FITC as a marker; ii) the cis-Golgi marker, GM130; or iii) the mitochondrial marker, Tom20. The tripartite combination induced distinct changes in the cellular distribution of the three markers (**Fig. 3A-C**). Importantly, the MakA/B/E tripartite disrupted the actin filaments (**Fig. 3A**) and induced Golgi fragmentation (**Fig. 3B**). The tripartite protein also caused rounding of the mitochondria, as evidenced by redistribution of the Tom20 staining from filamentous mitochondria to round structures (**Fig. 3C**). The individual components of the tripartite complex MakB or MakE failed to induce any detectable changes in the cellular distribution of markers discussed above. However, MakA had a weak effect on the disruption of Golgi and Mitochondria (**Supplementary Fig. 4A-C**). To investigate if rounding of the mitochondria led to mitochondrial dysfunction, HCT8 cells were treated with MakA or MakB or MakE or the MakA/B/E tripartite complex and stained for mitochondrial potential marker, TMRM. Similar to changes in the organelle structure, the tripartite MakA/B/E complex caused depolarization of mitochondria (**Fig. 3D-E**). The individual components of the complex had very little to no effect on the mitochondrial potential of the epithelial cells (**Fig. 3D-E**). In addition to depolarization of the mitochondrial membrane, the effect on total cellular ATP was measured after treatment with the individual and tripartite MakA/B/E proteins. The tripartite caused the most severe depletion of the total cellular ATP in the colon carcinoma (CaCO2) cells (**Fig. 3F**). Importantly, the tripartite caused a time-dependent decrease in the total cellular ATP content of the CaCO2 cells (**Fig. 3G**). Together, these results suggest that the Mak tripartite mediated disruption of cell organelles leading to dysfunction of the mitochondria.

**Figure 3.**
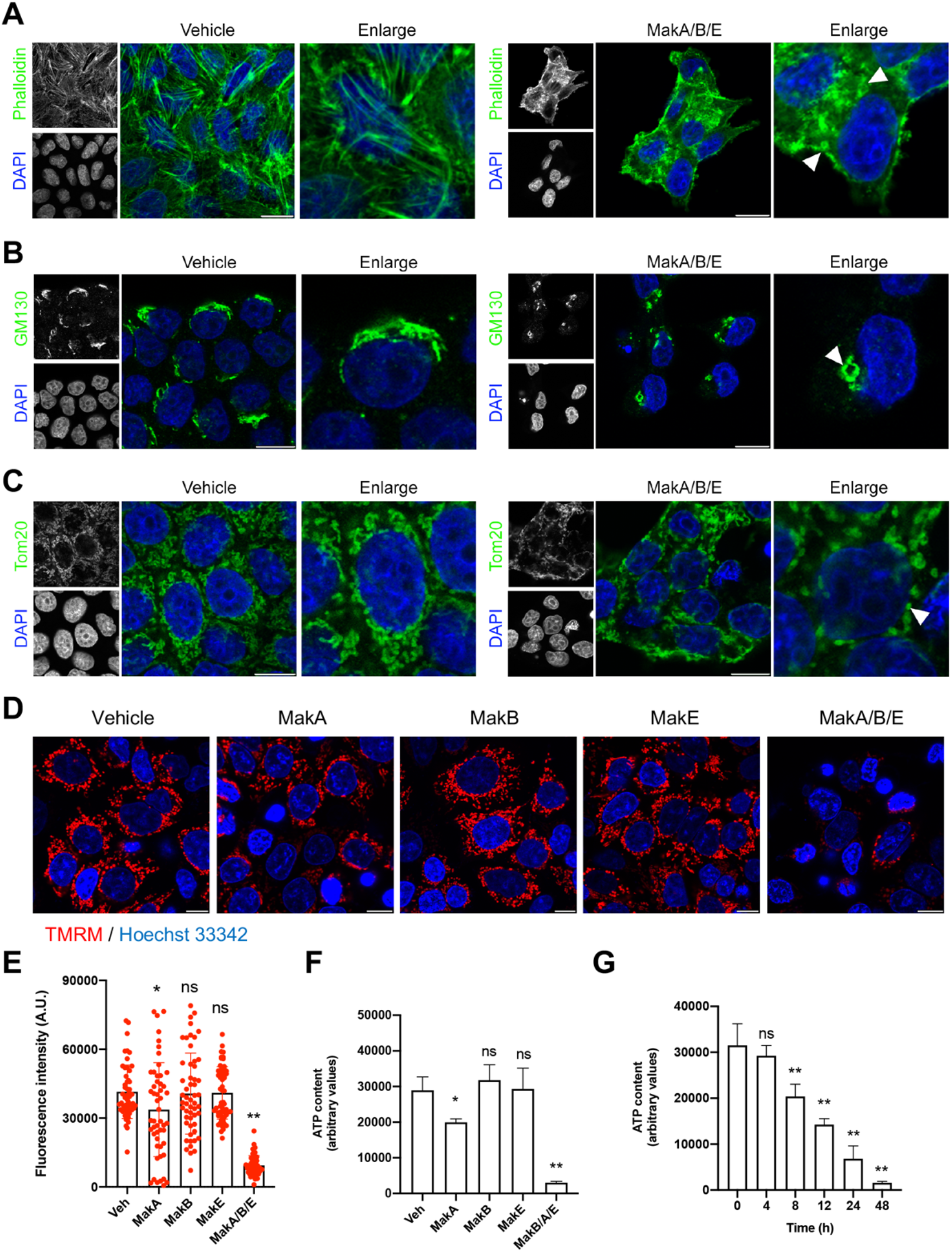
Effect of the tripartite Mak cytotoxin on intracellular structures and organelles. CaCO2 cells treated with Vehicle or MakA/B/E (250 nM, equimolar concentration) for 24 h were examined by confocal laser scanning microscopy. Nuclei were counterstained with DAPI. Scale bars represent 10 µm. ***(A)*** Effect on actin filaments. Cells were stained with Phalloidin-FITC to visualize actin filaments. Arrowheads (white) indicate disruption of actin filaments. ***(B)*** Effect on the Golgi apparatus. Cells were stained using a cis-Golgi marker, GM130. Arrowhead (white) indicates changes in the cellular distribution of the Golgi complexes. ***(C)*** Effect on mitochondria. Immunofluorescence detection was performed with antibodies against Tom20 to visualize mitochondria. Arrowhead (white) indicate swelling of mitochondria. ***(D)*** HCT8 cells treated were with a vehicle, the individual component of the tripartite complex or MakA/B/E (24 h) and stained with mitochondrial potential marker, TMRM (250nM, 30 min). Scale bars represent 10 µm. ***(E)*** Bar graphs show mean ± s.e.m. Data points represent fluorescence intensity of 51-57 individual cells. Significance was determined from the replicates using one-way analysis of variance (ANOVA) with Dunnett’s multiple comparisons test. *p<0.05, **p<0.01, ns = not significant. ***(F)*** Effect on cellular ATP content. CaCO2 cells were treated with 250 nM of the MakA, MakB and MakE proteins, individually and in the tripartite combination at equimolar concentration, for 48 h. Changes in cellular ATP contents were quantified as described in Supplementary information. Histogram represents data from four biologically independent experiments. Bar graphs show mean ± s.d. Significance was determined from the replicates using one-way analysis of variance (ANOVA) with Dunnett’s multiple comparisons test. *p<0.05, **p<0.01, ns = not significant. ***(G)*** CaCO2 cells were treated with equimolar concentration of the MakA/B/E tripartite (250 nM) in a time-dependent manner. Histogram represents data from three independent experiments. Bar graphs show mean ± s.d. Significance was determined from the replicates using one-way analysis of variance (ANOVA) with Dunnett’s multiple comparisons test. *p<0.05, **p<0.01, ns = not significant.

### Structural basis for a tripartite cytolytic toxin formed by secreted Mak proteins

MakA (PDB 6EZV) crystal structure provided the first molecular details of a protein encoded by the *mak* operon (14). MakA is organized in two domains, a long tail domain consisting of a five-helix bundle and a head domain comprising shorter helices and a β-sheet (strand order β1β3β2β4). MakB and MakE were predicted to have the same fold as MakA, although the low sequence identity (20% with MakB and 25% with MakE) made structure determination by molecular replacement less feasible. Therefore, selenomethionine (SeMet) labeled MakB and MakE were prepared and used for single-wavelength anomalous data diffraction (SAD) phasing. For both proteins, initial models were built from the SeMet-phased electron density maps and used for further model building using the native data. MakB and MakE were refined to 2.1 and 2.0 å resolution, respectively (**Supplementary Table 1)**. As in solution, their structures and crystal packing are monomeric with no obvious formation of oligomers.

MakE is structurally very similar to MakA, with a root mean square deviation of 2.3 å calculated on 329 aligned Cα atoms. MakE is a two-domain protein consisting of a tail and a head (**Fig. 4A**). The tail domain, 85 Å in length, consists of five long helices (helix 1, 2, 3, 6 and 7). The head domain, 55 Å in length, comprises two parallel helices (α4 and α5) and a four-stranded antiparallel β-sheet (strand order β1β5β2β6). β1 is very short and originates from the tail domain. MakE also has a β-hairpin (β3β4) protruding 90° from the sheet. Two molecules of MakE are found in the asymmetric unit, along with two nickel ions and four sulphate ions.

**Figure 4.**
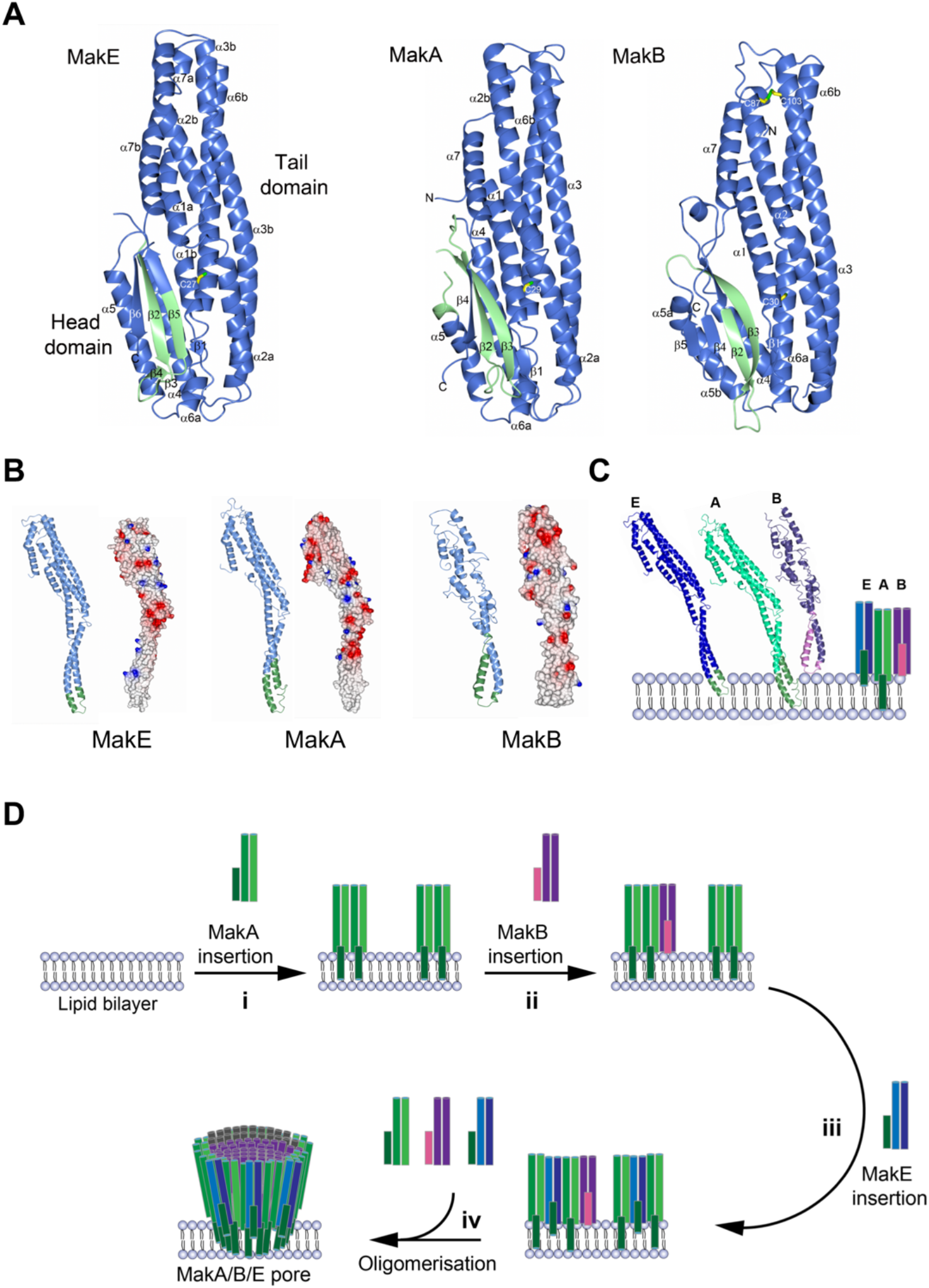
The crystal structures of MakE, MakA and MakB. ***(A)*** The crystal structures of MakE, MakA and MakB proteins are presented as ribbon diagrams, depicted in blue with the hydrophobic region (β-tongue) in green. The conserved cysteine (located on the first helix of all MakB/A/E proteins) and the disulfide in MakB are indicated as a stick models. The head and tail domains are labelled. ***(B)*** Elongated models of MakE, MakA, and MakE, calculated from the AhlB and AhlC structures (21, 51). The region originating from the β-tongue is shown in green. Their electrostatic properties are shown in the same orientation. The surface potential was calculated with ccp4mg (52). ***(C-D)*** A schematic illustration of how a combination of the Mak proteins may form a tripartite pore complex in a membrane. **i)** MakA is predicted to form two transmembrane helices, which enables MakA to anchor to the membrane. **ii)** MakB is less hydrophobic and binds to MakA. **iii)** The hydrophobic region of MakE is shorter and can only traverse one leaflet of the membrane. **iv)** In order to associate on the membrane, MakE would need to interact with the MakA/ complexMakB forms the scaffold to which MakA and MakE bind and a pore is formed. Their hydrophobic regions are shown in green (MakE and MakA) and pink (MakB).

MakB, encoded by the third gene of the operon, directly upstream of *makA*, has the same overall fold as MakA and MakE but deviates more from MakA with a root mean square deviation of 3.0 Å calculated on 328 Cα atoms. The MakB tail domain is very similar to that of MakA; the differences between the two proteins are mainly in the head domain in which MakB comprises a larger β-sheet (strand order β1β3β2β4β5) than MakA. One MakB molecule is found in the asymmetric unit in which we also modelled two sulphate ions. A comparison of MakA, MakB and MakE clearly revealed the similarity of their orientation and tight packing between the head and tail domains. In the tail domains, all three proteins have a bend in the α2 helix in equivalent positions. In MakA and MakB, the bend is formed in close proximity to a proline (Pro70 and Pro67, respectively). MakE, which has the most prominent break of the α2 helix, somewhat surprisingly has no proline at this position. All three proteins have a conserved cysteine positioned in the tail domain. In MakA and MakE, the cysteine is a few residues from the end of helix α1 (Cys29 and Cys27, respectively), and in MakB, it is the final residue of α1 (Cys30) (**Fig. 4A**). MakB has two additional cysteines, Cys87 and Cys103, which form a disulfide bond that ridgifies the connection between the C-terminus of α2 with the N-terminus of α3. The head domains share the same overall fold, but they are more structurally diverse from each other as they differ in the number of β-strands and in the length of the loops.

Except for the cysteines described above, amino acids sequence alignment revealed very few conserved residues. For instance, MakA has a FTPP motif located in the first strand, reminiscent of the Fxxxφ motif (where φ is any hydrophobic residue) needed for transport of flagellar rod and hook proteins. We have previously investigated if Phe37 of the motif is important for the delivery of MakA via the flagellum, and indeed the F37D mutant shows reduced secretion levels (14). However, we could not identify any similar motif in the corresponding region in MakE and MakB (**Supplementary Fig. 5**).

### Predicted transmembrane helices in MakA, MakB and MakE

The predictions program TMHMM (27) and Kyte and Doolittle hydropathy plots predicted transmembrane helices, or a hydrophobic region, in the head domain for all three proteins, MakA, MakB, and MakE (**Supplementary Fig. 5 and Supplementary Fig. 6**). In MakA the residues 198-246 were predicted to fold as two transmembrane helices, and in MakE one transmembrane helix was predicted from residues 202-224. In MakB, the program detected a stretch with increased hydrophobicity, residues 194-228, however a clear transmembrane helix was not suggested. When these regions are mapped onto the respective crystal structures, it is clear that they build up the two central β-strands and the beginning of helix α5 of MakA. In contrast, the hydrophobic region in MakE constitutes the first central β-strand (β2), the β-hairpin β3β4, and half the next β-strand (β5). In MakB, the hydrophobic region constitutes most of β2 and all β3 of the β-sheet in the head domain **(Fig. 4A)**. Most of these hydrophobic residues are shielded from solvent by packing against helix α1 of the tail domain and the helices of the head domain. In accordance with the nomenclature used for ClyA, which has a similar hydrophobic stretch, we propose that these residues are referred to as “β-tongue”.

Analyses of MakA, MakB, and MakE with DALI (28) identified a number of structurally related proteins of bacterial origin. The top hit structures with Z-scores of 20 or higher in the DALI search were AhlB, SmhA and SmhB (PDB 6GRK, 7A27 and 6ZZ5) from *A. hydrophila and S. marcescens* (21, 22), followed by two toxin components from *B. cereus*, HBL-B and NheA (PDB 2NRJ and 4K1P, respectively) (29, 30) and the nematicidal toxin Cry6A (PDB 5GHE) from *Bacillus thuringiensis* (31). Notably, Ahl, HBL and Nhe have been described as tripartite toxins (20, 21). Further down the list (Z-score 18) was an elongated form of AhlB (PDB 6GRJ) followed by the bi-component toxins YaxA, XaxA, and PaxB with Z-scores of 11-13, (PDB 6EK7, 6GY8, and 6EK4) (32, 33). A protomer form of ClyA (PDB 2WCD) (34), was found with a Z-score of 11. In addition to the DALI search, we performed protein homology searches for the MakA, MakB and MakE using the Phyre2 database (35). The MakA, MakB, and MakE structurally homologous bacterial toxins identified by Phyre2 were further analyzed for a phylogenetic relationship that revealed clustering of MakA, MakB and MakE together with NheA (**Supplementary Fig. 7**). Furthermore, most of these structurally related proteins had a hydrophobic cluster ranging from 150-300 amino acid residues, except the XaxB and YaxB that lacked this region (**Supplementary Fig. 6**).

The food poisoning bacterium *B. cereus* expresses at least two tripartite toxins, HBL and Nhe (20). HBL consists of HBL-B, HBL-L1 and HBL-L2, and the Nhe enterotoxin of NheA, NheB and NheC. Of these toxins, the crystal structures of HBL-B and NheA have been determined (29, 30). Both of them are structurally very similar to MakA, MakB and MakE as described above, with head and tail domains located in similar configurations. In addition, HBL-L1, HBL-L2, NheB, and NheC are predicted to have similar folds.

Recently, the crystal structures of the *A. hydrophila* proteins AhlB and AhlC, components of the tripartite toxin AhlABC, and of the *S. marcescens* proteins SmhA and SmhB, components of the tripartite SmhABC toxin, were reported (21, 22). The respective third components, AhlA and SmhC, were predicted to have structures similar to that of the other Ahl and Smh proteins. Most of these proteins have a predicted hydrophobic β-tongue, located in an equivalent position as MakA, MakB and MakE (**Fig. 4A**). Interestingly, in each of the currently described tripartite toxins (Mak, HBL, Nhe, Ahl and Smh), one of the components (MakA, HBL-L1, NheB, AhlB and SmhB) exhibits a rather prominent hydrophobic β-tongue, equivalent to two transmembrane helices. A shorter hydrophobic stretch, equivalent to one transmembrane helix, is found in MakE, NheC, AhlC and SmhC, whereas less pronounced hydrophobic regions are found in MakB and HBL-B. In NheA, HBL-L2, AhlA and SmhA, the TMHMHH program did not identify any significant hydrophobic regions.

The Ahl proteins were determined in alternative crystal forms, which allowed the identification of two major folds. One AhlB fold is very similar to that seen in our crystals of MakA, MakB and MakE (thus the high Z-score), where the head and tail domains are tightly packed against each other. Like the Mak proteins, AhlB has the same break in α2, due to the presence of a proline and a cysteine located at the end of α1 **(Supplementary Fig. 8**). In the other AhlB and AhlC forms, the head domain is rearranged into a helical extension of the tail helices, resulting in a 150 Å-long protein. This form is described as the elongated pore form, and the hydrophobic β-tongue identified by the TMHMM program is now found at the end of α3 and the beginning of α4. We used the pore forms of AhlB and AhlC (PDB 6GRJ and 6H2E) to model elongated forms of MakA, MakB and MakE **(Fig. 4B**). As a result, we obtained a model of MakA with two hydrophobic regions at the cusp of α3 and α4, long enough to span over two leaflets of a membrane (38 Å). Similarly, an elongated form of MakE was obtained based on the same structure, resulting in a model where the tip of the long α3/α4 helices is hydrophobic, albeit the hydrophobic residues are more concentrated on one side. The pore structure of AhlC was used to model MakB and generated a structure where the tip of the protein has one more hydrophobic side and one more polar side (**Fig. 4B**).

### Formation of tripartite Mak toxin oligomers

MakA, MakB, and MakE act as monomers during size exclusion chromatography, eluting as single peaks at 15.6 mL on a Superdex200 10/300 column. When MakA, MakB and MakE were mixed and incubated with the detergents n-Dodecyl β-D-maltoside (DDM) or cymal5 in the presence of lipid extracts (LE) an elution peak at 13.0 mL was obtained, indicating that an oligomeric complex was formed. However, when incubated with lauryl maltose neopentyl glycol (LMNG) and LE the elution volume was shifted to 12.1 mL, which indicates the formation of a higher oligomeric complex. Any oligomers of MakA, MakB or MakE in bipartite combinations, purified using the same protocol could not be detected. **(Supplementary Fig. 9)**.

In the study of the Ahl toxin, it was shown that a triple leucine to threonine mutation (α3 L156T, and α4 L160T, L161T) in the AhlC transmembrane region reduced the hemolytic activity of the AhlABC tripartite complex (36). As MakA has high structural similarity to both AhlB and AhlC, we tested the effect of similar mutations in its hydrophobic head domain. *E. coli* strains with plasmids expressing the wild type *makB*^*+*^*A*^*+*^*E*^*+*^ genes or mutant constructs with alterations in the *makA* gene were used for *in vivo* tests with the *C. elegans* killing assay. In comparison with the wild type comstruct, the *E. coli* derivative harboring p*makB*^*+*^*A*_*L193T,L246T*_*E*^+^ showed clearly reduced toxicity towards *C. elegans*. Furthermore, the *E. coli* derivative harboring p*makB*^*+*^*A*_*L193T*,***L207T***,***L213T***,*L246T*_*E*^+^ appeared completely attenuated in its toxicity towards *C. elegans* (**Fig. 1C**). The level of expression of the Mak proteins remained unaltered whereas the mutant clones showed reduced secretion of MakA. while secretion of MakB and MakE appeared essentially unchanged (**Fig. 1D**). tThese results show that the hydrophobic head domain of MakA plays an important role for the toxicity towards *C. elegans*, and for efficient secretion of the MakA protein.

Consistent with the prediction by the DALI server (28), our results suggest that the MakA, MakB, and MakE proteins most likely can form a tripartitie pore in membranes in a manner resembling the structures described for AhlABC from *A. hydrophila* (21). To further investigate the possible mechanism of pore assembly of the MakA/B/E tripartite complex in the presence of lipid membranes, we performed liposome pull-down experiments as described in Material and methods In short, the liposomes were incubated with the MakA/B/E tripartite protein mixture or with the individual MakA, MakB or MakE proteins followed by a cross-linking step and centrifugation to pull-down any cross-linked liposome-protein-complexes to be subjected to western immunoblot analysis (**Supplementary Fig. 10**). When incubated with MakA/B/E, several oligomeric formswere detected with the MakA antisera. The most pronounced were the monomeric and dimeric forms. Incubation with MakA alone resulted in a similar pattern.

With the MakB and MakE antisera, mainly larger oligomeric forms were detected, but also dimers and tetramers (MakE) and trimers (MakB) appeared. Incubation with MakE alone resulted in detection of monomers only, whereas with MakB alone no protein was detected suggesting that MakB was not efficiently binding to the liposomes used in the pull-down assay (**Supplementary Fig. 10**). We conclude from the results that MakA cound bind to the liposomes independently of other proteins and that it formed both dimers, tetramers and larger oligomers. It appeared that MakE bound rather weakly to liposomes but did not oligomerize. In case of MakB alone we did not detect any binding to the liposomes in this assay. (**Supplementary Fig. 10**). These findings indicate that binding of MakA to the liposomes facilitated interaction and subsequent oligomerization of MakB and MakE together with MakA. On the basis of the results from these liposome pull-down experiments and from the above described (**Fig. 1E and F**) hemolysis assays with different sequential orders and combinations of Mak proteins we propose a model for the MakA/B/E tripartite lytic membrane pore formation (**Fig. 4C-D**). In short, we assume that MakA, MakB and MakE can be released from the bacteria via the Vibrio flagellum secretion system in their elongated forms where the head domains have rearranged into two long parallel helices. In MakA, the hydrophobic β-tongue is located at the ends of these helices and can penetrate both leaflets of the membrane, anchoring MakA firmly. Next, MakB, binds to the membrane-anchored MakA and arranges as a scaffold for the pore-like complex. Finally, MakE joins the assembly and interacts with MakA and/or MakB. Additionally, its hydrophobic region is long enough to penetrate one leaflet of the membrane.

### Phylogenetic distribution of the *mak* gene cluster among *Vibrionaceae* strains

In a quest to further investigate the biological relevance of Mak proteins, we focused on their genetic determinants organized as the discrete gene cluster, *makDCBAE*, found in *V. cholerae* strain A1552 (**Fig. 1A**). By bioinformatic analyses of over a hundred completely sequenced genomes of *Vibrio spp*., we found that the complete *mak* gene cluster is specifically distributed among distinct strains of three *Vibrio* species—*V. cholerae, V. anguillarum* and *V. qinghaiensis* (**Supplementary Table 2**). The strains include different serogroups (O1, O139 and non-O1/O139) and both the classical and the El Tor biovars. Among them are representatives for several of the proposed six stages in the evolution of the current seventh pandemic of *V. cholerae* (37). The gene cluster was predominantly located on the second chromosome (Chr. 2) of the analysed strains and was present in *V. cholerae* with a single chromosome (**Fig. 5; Supplementary Table 2**). A few species-specific characteristics also emerged. In the case of *V. cholerae* strains, the *mak* gene cluster is predominantly located in the clockwise orientation of their chromosomes and is flanked by nearby genetic elements putatively coding for activities implicated in lateral gene transfer (LGT), integrases, and transposases (**Fig. 5; Supplementary Table 2**). The *mak* genes found on the genomes of *V. anguillarum* and *V. qinghaiensis* strains are mostly present in the counterclockwise orientation, and they are not immediately bordered by LGT-promoting genetic elements (**Fig. 5; Supplementary Table 2**).

**Figure 5.**
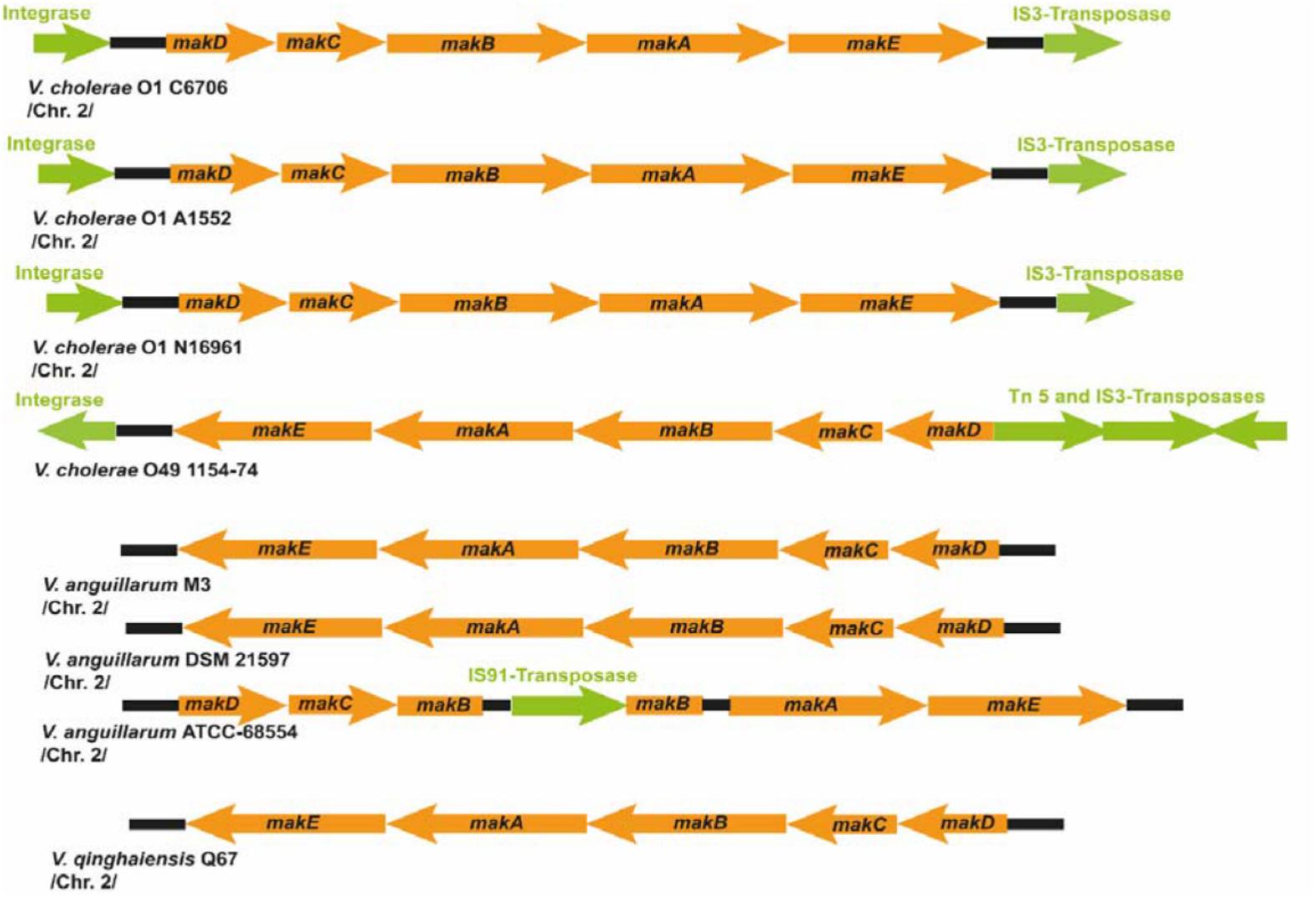
Presence of the *mak* gene cluster in different *Vibrionaceae* strains. Gene organization of *mak* gene clusters in the genomes of representative strains of *V. cholerae, V. anguillarum* and *V. qinghalensis*. Genetic information of transposable elements in flanking regions and within the gene clusters are indicated in green.

These findings, together with the estimation that *V. anguillarum* emerged earlier in evolution than *V. cholerae* (38), led us to explore the phylogenetic distribution of the *mak* gene cluster among the *Vibrionaceae* strains listed in **Supplementary Table 2**. We constructed a phylogenetic tree using the *mak* gene cluster of *V. cholerae* with the flanking LGT-promoting genes. The tree depicted two main groups: one composed of the strains of *V. anguillarum* and *V. qinghaiensis*, and the other group formed by the strains of *V. cholerae* (**Supplementary Fig. 11**). The calculated evolutionary distance additionally suggests that the *mak* operon was acquired earlier and then preserved by *V. anguillarum*, consistent with the fact that the *mak* genes of *V. anguillarum* strains lack the flanking the LGT genes (**Fig. 5; Supplementary Fig. 11**). In *V. cholerae*, the *mak* genes may have been acquired more recently. In addition to putative transposase and integrase genes, the lower GC content (41%) compared to the overall genomic GC content (around 47.5%) is consistent with the idea that the *mak* operon of *V. cholerae* strains may be an acquired genomic island.

## DISCUSSION

Our earlier identification and structural characterization of MakA (14), prompted this investigation on other proteins encoded by the same gene cluster. Here we establish that MakA, MakB and MakE, together can form a tripartite cytolytic toxin. The *V. cholerae makDCBAE* operon is subject to positive regulation by HapR, a transcriptional regulator of quorum sensing that also acts as a repressor for cholera toxin expression (14). In addition, transcription of the *makDCBAE* operon is growth phase-dependent; it is specifically activated in the stationary growth phases by the transcriptional regulator RpoS (39). The Mak proteins are thereby expected to be less expressed under conditions of acute cholera disease. Instead, they are more likely produced during bacterial growth in the natural aquatic environments where predators and nutrient limitations challenge bacterial fitness and survival. This is the first described tripartite cytotoxin from *V. cholerae* and our findings are important for the further understanding of its potential role in bacterial fitness and pathogenesis during infection in different hosts and environments.

For our structural analyses the MakA, MakB and MakE proteins were crystallized in their soluble, compact forms where a hydrophobic region, the β-tongue, is hidden in the interface between their respective head and tail domain. This region is longer in MakA (48 residues) than in MakE (22 residues), whereas the equivalent region in MakB is more amphipathic. In order to assemble into a membrane-bound tripartite complex, the important prerequisite is that each protein is predicted to undergo a conformational change where the head domain is unfolded and rearranged into two long helices. This model was discussed for the tripartite Ahl and Sml toxins (21, 22) but was comprehensively addressed in the case of the pore-forming toxin ClyA (40). Based on the above proposed theory, the tripartite assembly of MakA/B/E can be postulated as follows. In MakA, the hydrophobic β-tongue region that is located at the end of helices, is long enough to traverse the whole membrane. MakE, with its shorter β-tongue, may be able to penetrate only a single membrane leaflet. On the other hand, because MakB is less hydrophobic, it is considered to not bind to the membrane. Consequently, one can expect that MakA would anchor to the membrane due to its more pronounced hydrophobicity and thereafter attract MakB and MakE to the complex.

The function of MakA as the seed for the formation of the tripartite complex was further demonstrated by the results when MakA, MakB and MakE were added to erythrocytes in different sequential orders to study the assembly pathway. When all proteins were added together, or when the proteins were added in the order MakA -> MakB -> MakE with 15 minutes intervals, the level of hemolysis was as high as in the case of the positive control, the prototypic pore-forming toxin ClyA (**Fig. 1E**). Hemolysis was also observed, albeit at a reduced level, when MakA and MakB were added together, followed by the addition of MakE. In all other combinations, when erythrocytes were treated with MakB and/or MakE before the addition of MakA, no or very little hemolysis was observed. We hypothesize that the first step is the most crucial, the conformational change of MakA and its subsequent anchoring to the membrane. Membrane-bound MakA attracted MakB, which could undergo a similar conformational change and interact with MakA but not directly with the membrane. Next, MakE, after the structural rearrangement, joined with the MakA/B complex and with the outer leaflet of the membrane via the hydrophobic helices (**Figure 4D**). The primary structure and smaller size of the MakC and MakD proteins differ from the MakA/B/E proteins, and we concluded that MakC/D were not required for Mak cytotoxicity *per se*. We propose that they merely function as accessory proteins during MakA/B/E protein toxin biogenesis/secretion, but their actual role is yet to be discovered.

In addition to the hemolytic activity and the toxicity to *C. elegans*, we investigated the effect of MakA/B/E on colon carcinoma cells. Some cytotoxicity could be detected by treatment with MakA alone, probably due to its capacity to assemble on the membrane. The MakA/B/E complex was considerably more cytotoxic and resulted in destabilization of actin filaments and destruction of mitochondria and the Golgi apparatus. Taken together, these studies provided evidence that the tripartite MakA/B/E toxin complex has the capability to cause such effects if presented during host interactions. However, it can be questioned if these observations provide a bona fide reason to assume that for example mitochondrial disruption is a terminal, or even intended, outcome of MakA/B/E activity. Our *in vitro* results merely show that this can be the outcome of MakA/B/E activity in the mammalian cell.

Recent *in vitro* studies with the MakA protein have revealed that it can act as a toxin on its own as well, and upon internalization may cause induction of apoptotic responses and autophagy in cultured mammalian cells (41-43). It is an interesting possibility that the MakA protein can play a dual role as a toxin both by itself and by being part of the tripartite cytolytic complex as shown in the present study. Notably, some MakA secretion was observed also from the flagellum-deficient bacteria and the three Mak proteins appeared not to be dependent on each other *per se* for secretion. However, efficient secretion of all three Mak proteins was found to be dependent on the bacterial flagellum machinery, and their coordinated gene expression by virtue of the *mak* operon organization strongly suggests that these Mak proteins evolved to be primarily associated with the tripartite protein complex. The fact that *V. cholerae* normally has a single polar flagellum is particularly interesting in the context of the Mak toxin secretion. The single flagellum would ensure that secretion of the three toxin subunits may occur in a spatially coordinated fashion directly to a target membrane.

In addition to their structural similarities among the five different bacterial tripartite protein toxins that have been described so far there are interesting questions to be clarified regarding their biogenesis. HBL and Nhe are expressed by the Gram-positive bacterium *B. cereus*, and Ahl, Smh and Mak are expressed by the Gram-negative bacteria *A. hydrophila, S. marcescens* and *V. cholerae*, respectively. The HBL and Nhe proteins are produced with N-terminal signal sequences indicating that they are transported over the membrane *via* the *Sec* pathway (44). Apart from the information that secretion of the HBL protein requires functional flagella (45), the actual mechanism of secretion remains to be clarified and it is feasible to consider a route involving the flagella as shown for MakA. The Nhe proteins are presumably exported via an alternative mechanism that is not yet identified (46). It is not known how the Ahl proteins are exported, but *S. marcescens* has flagella. *e*.*g*. fT3SS (47) and *A. hydrophila* has both a vT3SS secretion system and flagella (48).

While the Mak proteins presumably play an important role for the fitness and survival of Vibrios in different hosts, the *makDCBAE* operon located on Chr. 2 may also be viewed as an example how an operon can be handled differently by evolution in various species of Vibrios that may have faced similar challenges. The similarities, such as LGT genes flanking the operon, the low G+C content (41%) and the presence of the degenerated *attB* site sequences upstream of the promoter proximal gene (*makD)*, that are shared with the *V. cholerae* pathogenicity islands VP1(49), and VSP-1 and VSP-2 (50), are in support of the notion that the *makDCBAE* operon represents a chromosomal genomic island. While the *makDCBAE* operon of *V. cholerae* may be considered in the process of assimilation, as evidenced by the remnants of the LGT genetic determinant, the *makDCBAE* gene cluster of *V. anguillarum* seems to be fully assimilated on Chr. 2 of this species. Our present findings will also be of particular relevance for further understanding of the *V. anguillarium* virulence and pathology during disease in fish, which is causing high mortalities and economic losses in aquaculture. We envision that the *mak* genes and proteins may be explored as novel potential targets for new diagnostics and therapeutic strategies in great demand towards the different pathogenic *Vibrionaceae*.

## MATERIALS AND METHODS

Details regarding cloning, protein expression and purification, assays, crystal structure determination, immunofluorescence, immunoblots and bioinformatic analyses are provided as **Supplementary Material**.

Data availability. Crystallographic coordinates and structure factors have been deposited in the Protein Data Bank under accession codes 6TAO (MakE) and 6T8D (MakB).

## Acknowledgements

This work was performed within Umeå Centre for Microbial Research (UCMR) and R.N. was supported from the UCMR postdoctoral programme, partially funded as a Linnaeus Centre of Excellence by The Swedish Research Council (2007-08673). K.P. received support from the Swedish Research Council (2016-05009) and The Kempe Foundations (SMK-1756.2 and SMK-1553). S.N.W. received support from the Swedish Research Council (2018-02914), The Kempe Foundations (JCK-1728), The Swedish Cancer Society (2017-419) and the faculty of Medicine at Umeå University. B.E.U. received support from the Swedish Research Council (2019-01720). We thank the Protein Expertise Platform (PEP) at Umeå University for help with construct design and cloning. We acknowledge the Umeå University Biochemical Imaging Center (BICU), Umeå Core Facility Electron Microscopy (UCEM) and the National Microscopy Infrastructure (NMI) for providing assistance in microscopy.

