## Supplementary material for "A tripartite cytolytic toxin formed by *Vibrio cholerae* proteins with flagellum-facilitated secretion"

#### Supplementary information

**Contents:**

Materials and Methods
Supporting Figures 1-11
Supporting Tables 1-4
References

**Material and Methods:**

**Bacterial strains and plasmids**

All bacterial strains and plasmids used in this study are listed in **Supplementary Table 3**.

***Vibrio cholerae* strain mutagenesis and growth conditions**

The *Vibrio cholerae* strains used for this study were the wild-type *V. cholerae* O1 El Tor strain A1552 and derivatives with the following mutations described earlier: *ΔmakA*, *ΔmakB*, *ΔmakC*, *ΔmakD* and *ΔflhA,* respectively (1). For the construction of a *ΔmakE* mutant derivative, the oligonucleotide primers used were based on *V. cholerae* O1 El Tor whole genome sequence (2). PCR amplification was performed using Phusion DNA polymerase (Thermo Scientific) and primers with the following sequences:

VCA0884A: CATTCTAGAAGCTGCAGTCTCCGGCGGTATC
VCA0884B CCCATCCACTATAAACTAACAAGAATAGAAGGGGGCGTAACC
VCA0884C TGTTAGTTTATAGTGGATGGGAAGTAATTGATTTCGCTCCAG
VCA0884D CTTCTAGACTGATGGGAAAGCTCATTAAAC

The *makE* deletion in *V. cholerae* was obtained using the procedure with positive selection for allelic exchange as described (3). For analyses of Mak protein expression and secretion, the *V. cholerae* strains were grown in LB media and incubated at 37 °C as previously described (1).

**MakB and MakC complementation experiments**

The *makB* and *makC* clones in the pBAD18 vector (1) were electroporated into *ΔmakB* and *ΔmakC* mutants, respectively. As a negative control, the expression vectors pBAD18 without an insert were introduced into the mutant strains.

**Cloning, overexpression and purification of Mak proteins.**

The details about MakA cloning, overexpression and purification were described earlier (1)The *makB* and *makE* genes (UniProt accession codes Q9KL65 and Q9KL63) were PCR amplified from genomic DNA of *V. cholerae* strain A1552. The *makB* PCR product was digested with *NcoI/XhoI*, the *makE* with *BspHI/HindIII*. Each product was subsequently cloned into equivalent sites of pET-His1a, in-frame with a cleavable histidine affinity tag with the sequence MKHHHHHHPMSDYDIPTTENLYFQGA. The plasmids were introduced by transformation into *E. coli* DH5α and positive colonies were subsequently sequenced. The constructs were expressed in *E. coli* BL21(DE3) in LB broth supplemented with 50 µg/mL kanamycin. When the cultures reached $OD_{600}$ ~0.6, protein expression was induced with 0.5 mM isopropyl 1-thio-β-d-galactopyranoside (IPTG) followed by growth for 5 h at 25 °C. Cells were harvested by centrifugation, and the pellets were stored at -80 °C until further use. The cell pellets were resuspended in 50 mM Tris-HCl pH 7.6, 0.3M NaCl, and 10 mM imidazole (lysis buffer) supplemented with 1% triton-X100 and sonicated on ice. The lysate was centrifuged at 63,000 x *g* for 20 min. The resulting supernatant was passed over a column packed with His60 Ni-resin (Takara). The column was washed with lysis buffer containing 30 mM imidazole after which the protein was eluted with the same buffer containing 0.3 M imidazole. Next, the histidine tag was removed by incubation with 1% (w/w) TEV protease overnight at 4 °C. After buffer exchange to 50 mM Tris-HCl pH 7.6, 0.2 M NaCl the protein solution was passed over the affinity column again. The flowthrough and wash fractions were collected and concentrated. The cleaved protein was further purified on a HiLoadTM 16/60 Superdex™200 prep-grade column (GE Healthcare) equilibrated with 20 mM Tris-HCl pH 7.6, 0.2 M NaCl. Fractions containing the peak of interest were concentrated in 20 mM Tris-HCl pH 7.6.

Selenomethionine (SeMet)-labelled MakB and MakE were produced by growing the cultures in M9 media supplemented with glucose at 37 °C. At OD600 ~0.4, 100 mg/L each of lysine, threonine, phenylalanine and 50 mg/L each of leucine, isoleucine, valine, proline and SeMet were added and protein expression was induced with 0.5 mM IPTG (4) at 20 °C overnight. The SeMet-labelled proteins were purified as described above.

**Crystallization and structure determination**

Sitting-drop vapor-diffusion trials were performed at 20 °C for MakB (8.3 mg/mL) and MakE (20 mg/mL) in 96-well MRC-crystallization plates (Molecular Dimensions). Droplets of 0.5 µL protein were mixed with equal volumes of screening solutions from Molecular Dimensions. For native MakB, the final optimized crystallization condition was 1.5 M $MgSO_4$ and 0.1 M MES pH 6.5. Crystals of SeMet-labelled MakB were obtained in 0.6 M ammonium sulfate, 0.1 M sodium acetate pH 4.5. Native MakE was crystallized in 10 mM $NiCl_2$, 0.1 M Tris-HCl pH 8.5 and 20% (w/v) monomethyl ether polyethylene glycol (PEG) 2000. SeMet-labelled MakE was crystallized in 24% (w/v) PEG 3350, 25 mM Na/K phosphate, 0.1 M BisTris propane pH 6.5.

#### Supplementary information

All crystals were soaked for 30 seconds in mother liquor solution supplemented with 20% (v/v) ethylene glycol or 20% (v/v) PEG 400 before they were flash cooled in liquid nitrogen and stored until data collection.

MakB native diffraction data were collected on a Pilatus3 2M detector at beamline ID23-2 and single anomalous diffraction (SAD) data at beamline ID23-1 on a Pilatus 6M F detector at the European Synchrotron Radiation Facility, Grenoble, France (ESRF). Native MakE data were collected at the ESRF beamline ID23-2 on a Pilatus3 2M detector. SAD data were collected at beamline BioMAX, MAX IV in Lund, Sweden. Diffraction images were processed with XDS (5) and scaled with Aimless (6) from the CCP4 program suite (7). The structure of SeMet-labelled MakB was solved with SAD-phasing using AutoRickshaw (8). Density modification and automatic model building were performed using AutoRickshaw and ArpWarp (9) and refined using phenix.refine (10). The SAD MakE data were carefully reprocessed using XDS (5), after which the scaled data were used as input in the CRANK2 pipeline (11). A search for ten Se atoms resulted in an initial model that was further built in COOT (12).

The crystals of the native MakE protein grew in space groups different from the SeMet-labelled proteins, and hence the structure was obtained by molecular replacement using Molrep (13, 14) with the SeMet structure as a search model. The native structures were refined using phenix.refine (10) and built using rounds of manual building in COOT (12). For the refinement of MakE, translational-libration-screw refinement was used, treating each molecule as an individual TLS group (15).

Figures were prepared with CCP4mg (16). Relevant processing and refinement statistics are summarized in Supplementary Table1. Elongated models of MakE, MakA, and MakB were obtained using Swissmodel (17) using the structures of AhlB (PDB 6GRJ) and AhlC (PDB 6H2D) as starting models(18).

**Analysis of MakA/B/E tripartite and bipartite oligomers using gel filtration**

The MakA/B/E tripartite were prepared by incubating equimolar concentrations (250 nM) of MakA, MakB and MakE with 0.1% detergent (Antrace) and lipid extracts (LE; Avanti Polar Lipids) at 25 °C for 1 hr. Subsequently, aggregates and MakA/B/E oligomers with high molecular weight were separated by size exclusion chromatography performed on Superdex 200 10/300 column (GE Healthcare) in 30 mM MES pH 5.6, 150 mM NaCl, 0.003% detergent and LE buffer. The detergents used were N-dodecyl b-C-maltoside (DDM), Lauryl Maltose Neopentyl Glycol (LMNG) and Cymal5. The bipartite combinations (250 nM) were purified using the same protocol.

#### Supplementary information

##### Antibodies

For immunodection of Mak proteins by immunofluorescence (IF) or western immunoblot (WB) analyses we used polyclonal antisera from rabbits (produced by GeneCust) at the following dilutions: Anti-MakA (IF= 1:100, WB= 1:5000), Anti-MakB (WB= 1:5000), Anti-MakC (WB= 1:5000), Anti-MakE (WB= 1:5000). Anti-Tom20 (#612278, IF = 1:100) and Anti-GM130 (#610822, IF = 1:100), mouse monoclonal antibodies were purchased from BD Biosciences. Anti-beta-actin mouse monoclonal antibody (#A2228, WB: 1:5000) was purchased from Sigma-Aldrich. Goat anti-Rabbit HRP-conjugated IgG (#AS09602, WB: 1:5000) were purchased from Agrisera AB, Sweden. Rabbit anti-mouse HRP-conjugated immunoglobulins (#P0260, WB: 1:5000) were purchased from Dako. Alexa Fluor 488/555/647 conjugated secondary antibodies for immunofluorescence were purchased from Thermo Fisher.

##### SDS-PAGE and Immunoblot analysis.

SDS-PAGE and immunoblot analysis were performed according to standard procedures(19). Briefly, bacterial whole cells and culture supernatants were separated by centrifugation at 10,000 × *g* for 15 min. Cell pellets were suspended in an appropriate volume of 1× SDS buffer and boiled to obtain whole cell lysates. Culture supernatants were filtered through a 0.45 μm PVDF syringe filter (Millipore, USA) and subsequently precipitated with 10% (w/v) trichloroacetic acid (TCA). The samples were centrifuged at 15,000 × *g* for 15 min at 4 °C to pellet the TCA-precipitated proteins, which were then resuspended in 1× SDS buffer and boiled for 5 min. Protein samples were resolved by SDS-PAGE and processed for immunoblotting. HRP-conjugated goat anti-rabbit IgG (Agrisera AB, Sweden) was used as a secondary antibody. Immunoblot detection was done using Clarity Western ECL substrate (BioRad). Prestained protein molecular weight standards (SM0679, Fermentas) were used to determine the protein molecular weights.

##### Hemolysis assay

For tests of hemolytic activity, Mak protein samples (250 nM) were mixed with a suspension of horse erythrocytes (2% of whole blood) in PBS and incubated 120 min at 37 °C. The cytolytic *E. coli* protein ClyA (250 nM) and Triton X-100 were were used as positive controls and PBS was the negative control. After centrifugation, the supernatants were monitored spectrophotometrically for released haemoglobin, by measurement of absorbance at 545 nm, as an indicator of red blood cell lysis. The ClyA protein was obtained as the native form from *E. coli* through the procedures for overproduction and purification described earlier (20, 21).

##### Cell lines and cell culture

#### Supplementary information

Human colon cancer cells (CaCO2; RRID:CVCL_0025 and HCT8; RRID:CVCL_2478) were obtained from American Type Culture Collection (ATCC) and maintained in RPMI-1640 or DMEM media supplemented with non-essential amino acids (1:100), sodium pyruvate (1 mM), penicillin (20 Units), streptomycin (20 µg/mL) and 10% fetal bovine serum (FBS) at 37 °C, 5% $CO_2$. All experiments were performed with mycoplasma-free cells.

**Cell viability and ATP estimation assays**

For cell viability experiments, CaCO2 or HCT8 colon carcinoma cells were grown on a 96-well plate ($5x10^3$/well, Tecan Group Ltd) overnight and treated with the individual Mak proteins for 48 h at 37 °C, 5% $CO_2$. Loss of cell viability was quantified by a decrease in MTS (Promega) absorbance. MTS absorbance (490 nm) was measured on an Infinite M200 microplate reader (Tecan) according to the manufacturer's instructions. Data were normalized to an equal volume of vehicle-treated (20 mM Tris-HCl, pH 7.4) cells and expressed as percentages.

For measurement of total cellular ATP content, CaCO2 cells were grown on a 96-well plate ($5x10^3$/well, Tecan Group Ltd) overnight and treated with increasing concentrations of the individual Mak proteins or the tripartite complex for 48 h or with the tripartite complex in a time-dependent manner at 37 °C, 5% $CO_2$. Cellular ATP content was measured with an ATPLite kit (PerkinElmer) on an Infinite M200 microplate reader (Tecan) according to the manufacturer's instructions.

**Immunofluorescence**

For fixed cell immunofluorescence, CaCO2 cells were grown in a coverslip-bottom 8-well chamber slide (µ-Slide, ibidi) and fixed in 4% paraformaldehyde for 30 min at room temperature (RT), permeabilized in 0.25% Triton X-100 (15 min). Cells were then washed with PBS and incubated with primary antibodies (overnight, 4 °C), followed by incubation with respective Alexa488/555 conjugated secondary antibodies. Nuclei were counterstained with DAPI (5 min, RT). For measurement of the mitochondrial potential, HCT8 cells were treated with an equal volume of vehicle (20mM Tris-HCl, pH 7.4), MakA, MakB, MakE or the tripartie complex (MakA/B/E) for 24 h. At the end of the treated cells were incubated with Image-iT™ TMRM reagent (Thermofisher) for 30 min (200 nM). Cells were visualized using a Leica SP8 inverted confocal system (Leica Microsystems) equipped with a HC PL APO 63x/1.40 oil immersion lens. Images were captured and processed using the LasX (Leica Microsystems). Images were processed using ImageJ – FIJI distribution (NIH) (22).

**Flow cytometry**

#### Supplementary information

Cellular uptake of Alexa568-MakA, Alexa568-MakB, Alexa568-MakE (250 nM, 24 h) or Alexa-MakA/B/E (Equimolar concentration of individual protein, 250 nM, 24 h) treated CaCO2 cells was investigated by flow cytometry (BD Accuri C6). Live cells were gated, and cellular uptake was represented as mean fluorescence intensity (MFI) (23).

**Lipids**

All lipids were purchased from Avanti Polar Lipids, Alabaster, AL, USA. Lipids: 1-palmitoyl-2-oleoyl-glycero-3-phosphocholine (POPS), 1,2-dioleoyl-sn-glycero-3-phosphoethanolamine (DOPE), 1-palmitoyl-2-oleoyl-sn-glycero-3-phospho-L-serine (POPS), Sphingomyelin from Porcine Brain (SM), Cholesterol from Ovine (Chol), L-α-phosphatidylinositol-4,5-bisphosphate from Porcine Brain (PIP2), and N-palmitoyl-sphingosine-1-{succinyl[methoxy(polyethylene glycol)5000]} (PEG5Kce). Lyophilized PIP2 lipids were dissolved in a mixture of Chloroform:Methanol (2:1) to a concentration of 1 mg/mL. Next, they were protonated by adding 0.5 µL of 1M HCl to 100 µg of PIP2, kept at room temperature for 15 min and dried down with nitrogen gas. The dried lipid was redissolved in a choloform:methanol (3:1) mixture to 1 mg/mL followed by drying again. Finally, the 100 µg of PIP2 was redissolved in 100% chloroform to 1 mg/mL and stored at -20 °C until used for liposome production.

**Liposome Preparation**

Liposomes containing 0.5 mol% PEG5Kce, 5 mol% PIP2, 10 mol% SM, 10 mol% POPS, 15 mol% DOPE, 20 mol% Chol, and 39.5 mol% POPC (referred herein to as synthetic lipid mixture, SLM liposomes) were prepared using the lipid film hydration and extrusion method. The SLM composition was inspired by the distribution of lipids found in the plasma membrane of HeLa cells (24) with slight adjustments made to the mole percentages to improve supported lipid bilayer production. The individual lipids dissolved in chloroform were mixed together, dried under nitrogen flow followed by a minimum of one hour. The dried lipid film was then rehydrated using a pre-heated citrate buffer at pH 4.5 (20 mM Citrate, 50 mM NaCl, pH 4.5, 37 °C) to a lipid concentration of 1 mg/mL. The solution was then extruded at ~40 °C eleven times through a polycarbonate membrane with 50 nm pore size using an Avanti miniextruder. The liposomes were stored at 4 °C until used.

**QCM-D**

The QCM-D measurements were carried out using an AWSensors X4 (AWsensors, Spain) instrument equipped with flow chambers and SiO2 coated sensors (Wrapped 14mm, 5 MHz, Cr/Au - SiO2, Polished). Each sensor was stored in 2% SDS overnight and treated with UV-ozone (Bioforce Nanosciences, USA) for 30 min prior to use. Supported lipid bilayers were formed using PM+PEG liposomes (100uL of 0.1 mg/mL) in citrate buffer at pH 4.5 (20 mM

Citrate, 50 mM, 0.1 mM EDTA, pH 4.5) followed by rinsing in the same buffer. The buffer was then exchanged for a buffer containing 120 mM citrate (pH 7), which was also the buffer used for protein adsorption. MakA, MakB and MakE (17.3 µM each) were mixed together in 120 mM citrate buffer for 10 minutes before being added. The flow was stopped once the protein reached the chamber, and the bilayer was incubated under steady conditions for 80 min before rinsing with the same buffer. The whole measurement was carried out at 37 °C. The protein layer was found to be rigid (D~$0.3*10^{-6}$). The frequency shift of the third overtone was therefore converted into a bound mass using the Saurbrey equation.

##### Liposome pull-down assay

*E. coli* total lipid extract (Avantis) dissolved in chloroform was dried to a thin film under nitrogen stream. The lipid film was allowed to hydrate in HEPES buffer saline (10 mM HEPES, 150 mM NaCl, pH 7.4). This suspension was extruded over polycarbonate membranes with 0.1 µm pore size using the Avanti Mini-Extruder (Avanti Polar Lipids, Alabaster, AL, USA). The liposomes (5 mg/mL) were incubated with MakA (500 nM), MakB (500 nM), MakE (500 nM) or the MakA/B/E tripartite complex (1:1:1) at 37 °C for 2 hours. At the end of the experiment, the reaction mixture was crosslinked with Glutaraldehyde (0.05 %) for 10 min at 37 °C followed by the addition of stop solution Tris-HCl (200 mM, pH 6.8). Subsequently, the reaction mixture was centrifuged at 21,500 x *g* for 30 mins. Liposome bound complexes were washed twice with the reaction buffer and loaded to the gel in the SDS-PAGE loading buffer. Liposome bound Mak proteins were detected using Mak proteins specific antibodies.

##### Transmission electron microscopy

For liposomes, negative staining samples were added to glow discharged 300 mesh copper grids with a thin film of carbon (Ted Pella, Redding, CA) washed twice with MQ water and negatively stained with 1.5% uranyl acetate solution (EMS (Hatfield, PA). Grids were examined with Talos L120C (FEI, Eindhoven, The Netherlands ) operating at 120kV. Micrographs were acquired with a Ceta 16M CCD camera (FEI, Eindhoven, The Netherlands) using TEM Image & Analysis software ver. 4.17 (FEI, Eindhoven, The Netherlands).

##### Analysis of the *mak* gene clusters in different *Vibrio* genomes

Sequences of the *mak* operons were retrieved from GenBank (E-value =0), based on searching with the *mak* gene cluster of *V. cholerae* O1 biovar El Tor str. N16961 chromosome 2 (accession number NC_002506) together with their flanking ORFs against the fully sequenced bacterial genomes using BLASTN. Sequences were aligned using the MUSCLE alignment algorithm, and the phylogenetic relationships among different *mak* gene clusters in the retrieved dataset were estimated based on the Maximum Likelihood (ML) method implemented in MEGA-X (25). The ML model of gamma distribution with invariant sites (G+I) was applied. The test of the clades was evaluated by 500 bootstrap replicates of Nearest-Neighbor-Interchange ML heuristic method.

#### Supplementary information

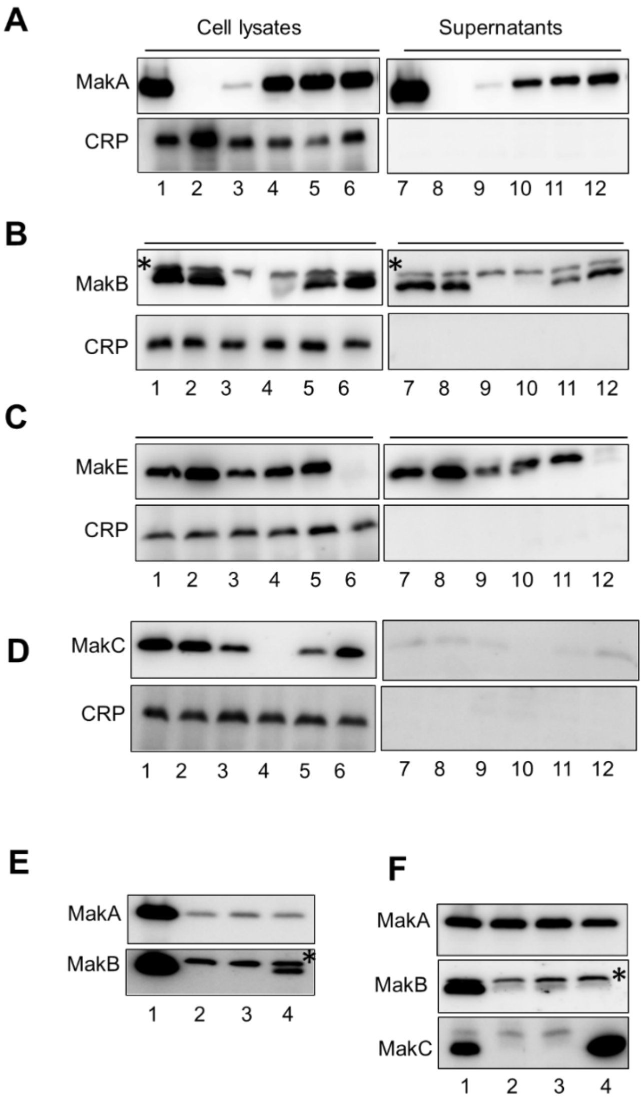

**Supplementary Fig. 1: *Vibrio cholerae* secretion of Mak proteins.**

***(A-D)*** Western immunoblot detection of Mak protein expression and secretion from *V. cholerae* O1 El Tor strain A1552. Samples of bacterial cell lysates (lanes 1-6) and culture supernatants (lanes 7-12) were subjected to immunodetection with antisera raised against MakA (A), MakB (B), MakE (C), and MakC (D) respectively. Lanes 1 and 7, wild type A1552; lanes 2 and 8, Δ*makA*; lanes 3 and 9, Δ*makB*; lanes 4 and 10, Δ*makC*; lanes 5 and 11, Δ*makD;* lanes 6 and 12, Δ*makE.* An additional band, marked with an asterisk, representing an unknown protein appeared in the case of the anti-MakB immunoblots. The cytoplasmic protein CRP was used as a reference and non-secreted protein control. Data are representative of three independent experiments. ***(E)*** Western blot analysis of whole cell lysates for MakA and MakB expression. Lane 1, wild-type A1552, lane 2, Δ*makB*; lane 3, Δ*makB*/pvector, lane 4, Δ*makB*/p*makB.* The asterisk indicates a nonspecific band detected by the anti-MakB antiserum. ***(F)*** Western blot analysis of MakA, MakB and MakC expression. Lane 1, wild-type A1552; lane 2, *ΔmakC*; lane 3, *ΔmakC/*pvector; lane 4, *ΔmakC/*pmakC. The asterisk indicates a nonspecific band detected by the anti-MakB antiserum.

#### Supplementary information

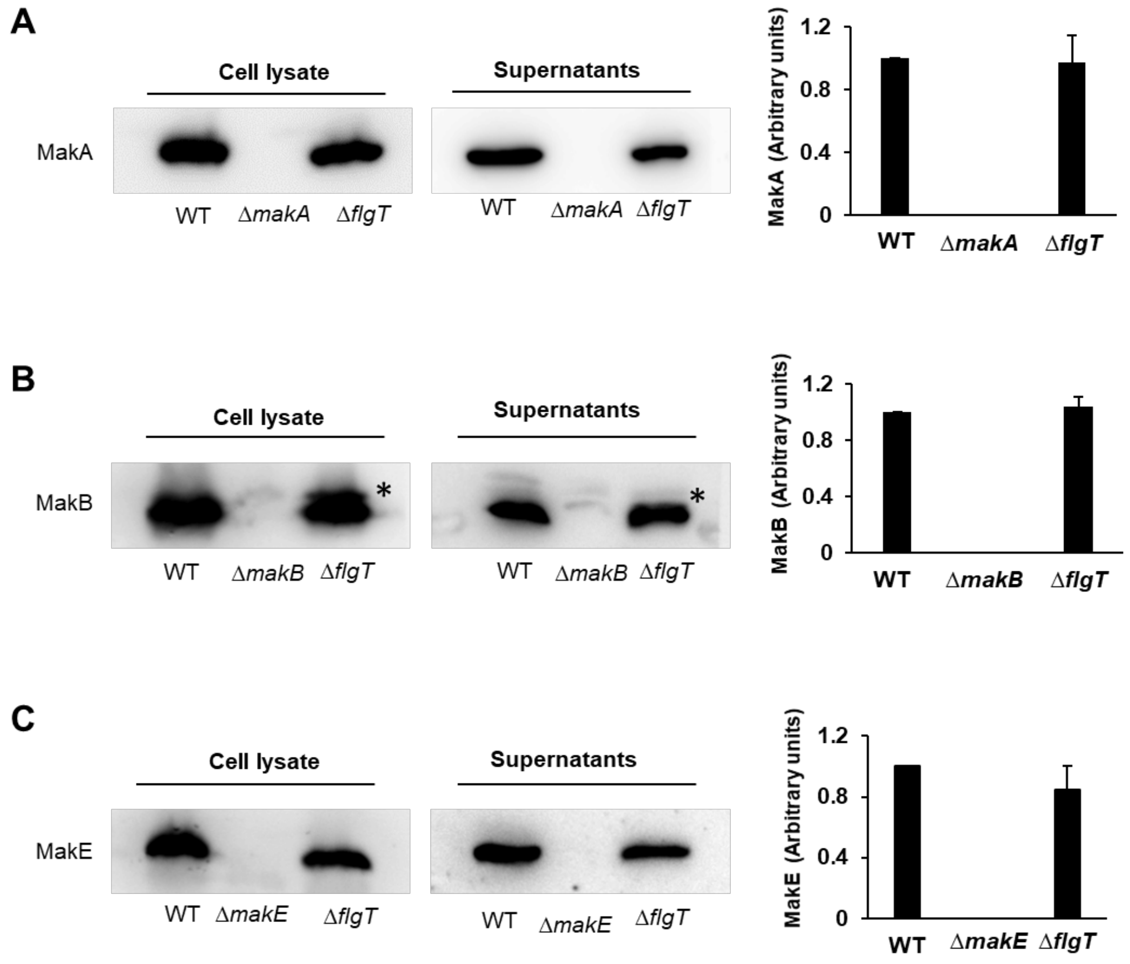

**Supplementary Fig. 2: *Vibrio cholerae* expression and secretion of the MakA/B/E proteins are not affected by lack of FlgT.**
Western blot analysis of Mak protein expression and secretion from *V. cholerae* O1 El Tor strain A1552 and Δ*flgT*. Samples of bacterial cell lysates or culture supernatants were subjected to immunodetection with antisera raised against MakA (**A**), MakB (**B**) and MakE (**C**), respectively. Samples from the respective Δ*mak* gene mutant derivatives of *V. cholerae* O1 El Tor strain are shown in the middle lane. Histograms to the right represent quantification of the Mak proteins. In (**B**), the asterisk indicates an unidentified protein also detected by the MakB antiserum. Data are representative of two independent experiments.

#### Supplementary information

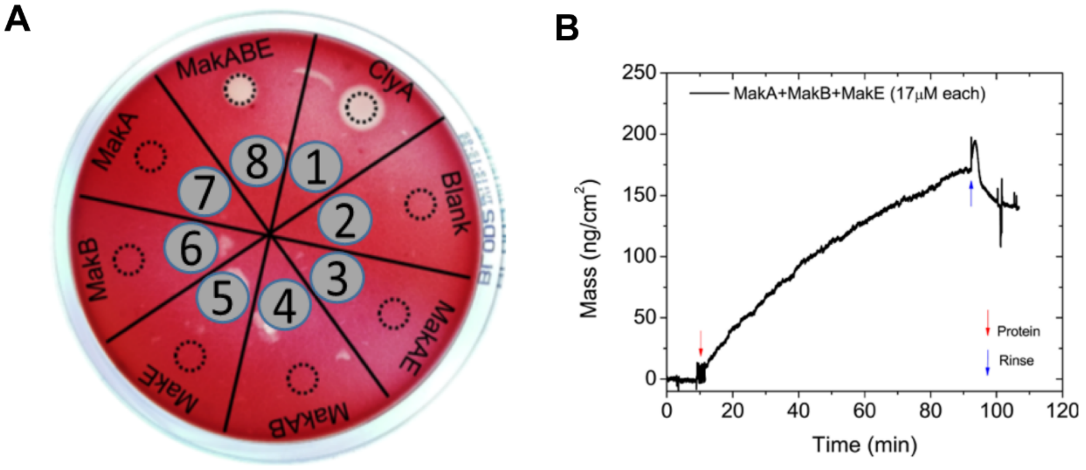

**Supplementary Fig. 3: The tripartite MakA/B/E cytotoxin binds to red blood cell and induces hemolysis**
***(A)*** Test of hemolysis activity of Mak proteins on blood agar. Samples (5 µl) of protein solutions were spotted onto an agar plate containing horse erythrocytes. The plate was incubated overnight at 37 °C, and a zone of clearance indicated lysis of the erythrocytes. The spots (marked by a dotted ring) were: 1) The cytolytic *E. coli* protein ClyA (100 nM) used as positive control; 2) No protein; 3) MakA+MakE (250 nM each); 4) MakA+MakB (250nM each); 5) MakE (250nM); 6) MakB (250 nM); 7) MakA (250nM); 8) MakA+MakB+MakE (250 nM each).
***(B)*** Quartz Crystal Microbalance with dissipation (QCM-D) data confirming the binding of a mixture of MakA, MakB and MakE (17 µM each) to a supported lipid bilayer made from the synthetic lipid mixture (SLM). The frequency shift of the third overtone was converted into a mass value using the Sauerbrey equation. The addition of the protein is marked with a red downwards arrow and onset of rinsing with a buffer with a blue upwards arrow.

#### Supplementary information

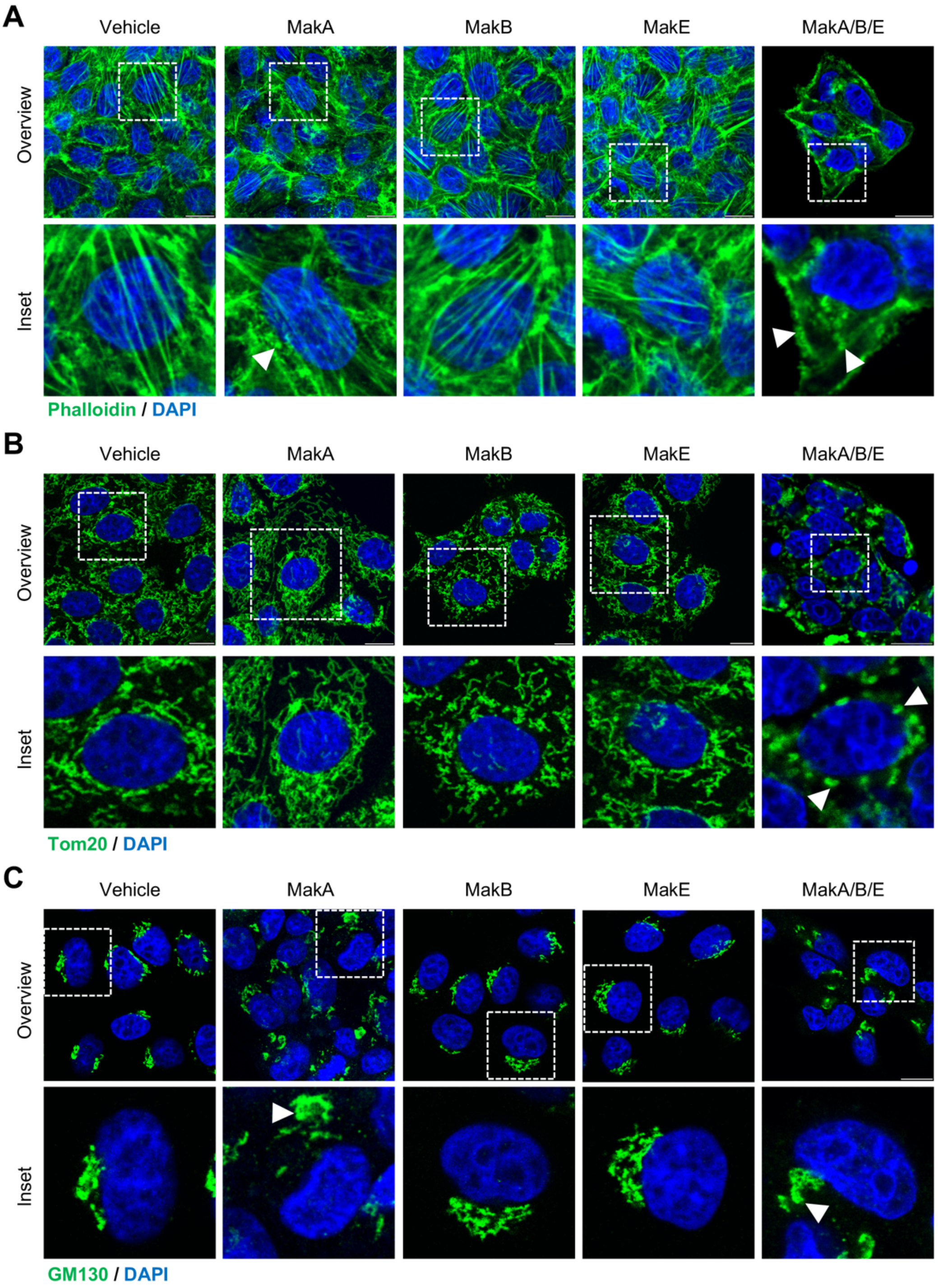

**Supplementary Fig. 4: Effect of the tripartite MakA/B/E cytotoxin or its individual component, MakA, MakB, or MakE, on intracellular structures and organelles.**
CaCO2 cells treated with an equal volume of vehicle, MakA (250 nM), MakB (250 nM), MakE (250 nM) or MakA/B/E cytotoxin (250 nM, equimolar concentration) for 24 h were examined by confocal laser scanning microscopy. Nuclei were counterstained with DAPI. Scale bars represent 10 μm.
***(A)*** Effect on actin filaments. Cells were stained with Phalloidin-FITC to visualize actin filaments. Arrowheads (white) indicate disruption of actin filaments.

***(B)*** Effect on the Golgi apparatus. Cells were stained using a cis-Golgi marker, GM130. Arrowhead (white) indicates changes in the cellular distribution of the Golgi complexes.
***(C)*** Effect on mitochondria. Immunofluorescence detection was performed with antibodies against Tom20 to visualize mitochondria. Arrowhead (white) indicate swelling of mitochondria.

#### Supplementary information

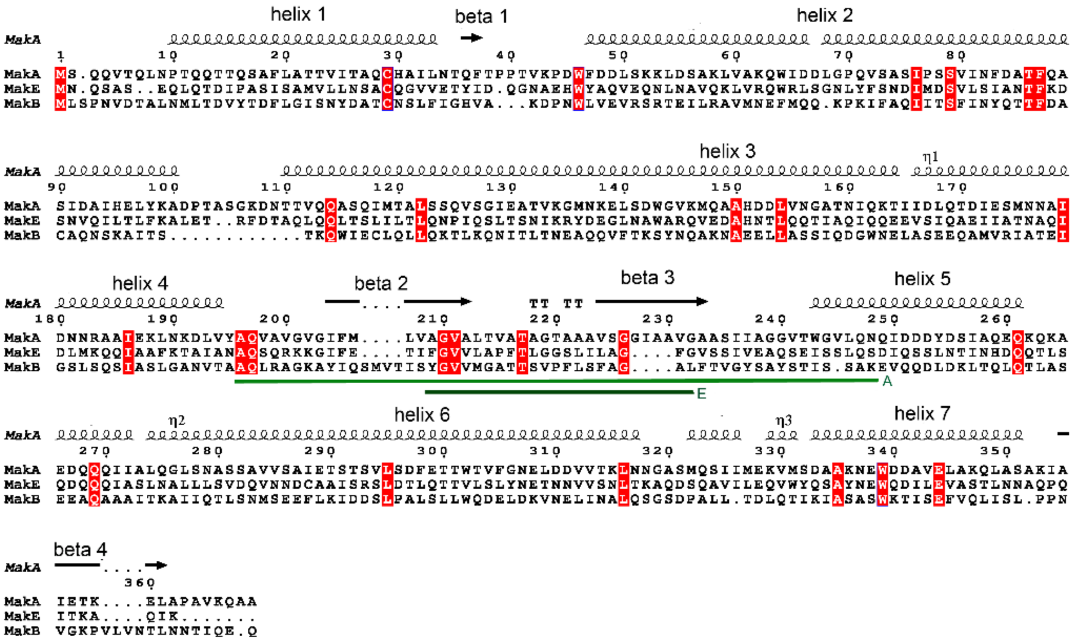

**Supplementary Fig. 5: Amino acids sequence comparison between MakA, MakE and MakB.**

Identical residues are highlighted in red. Secondary structure elements, based on the MakA structure, are indicated above the sequences. The residues in MakA and MakE predicted to be transmembrane helices are indicated with green bars below the sequence.

#### Supplementary information

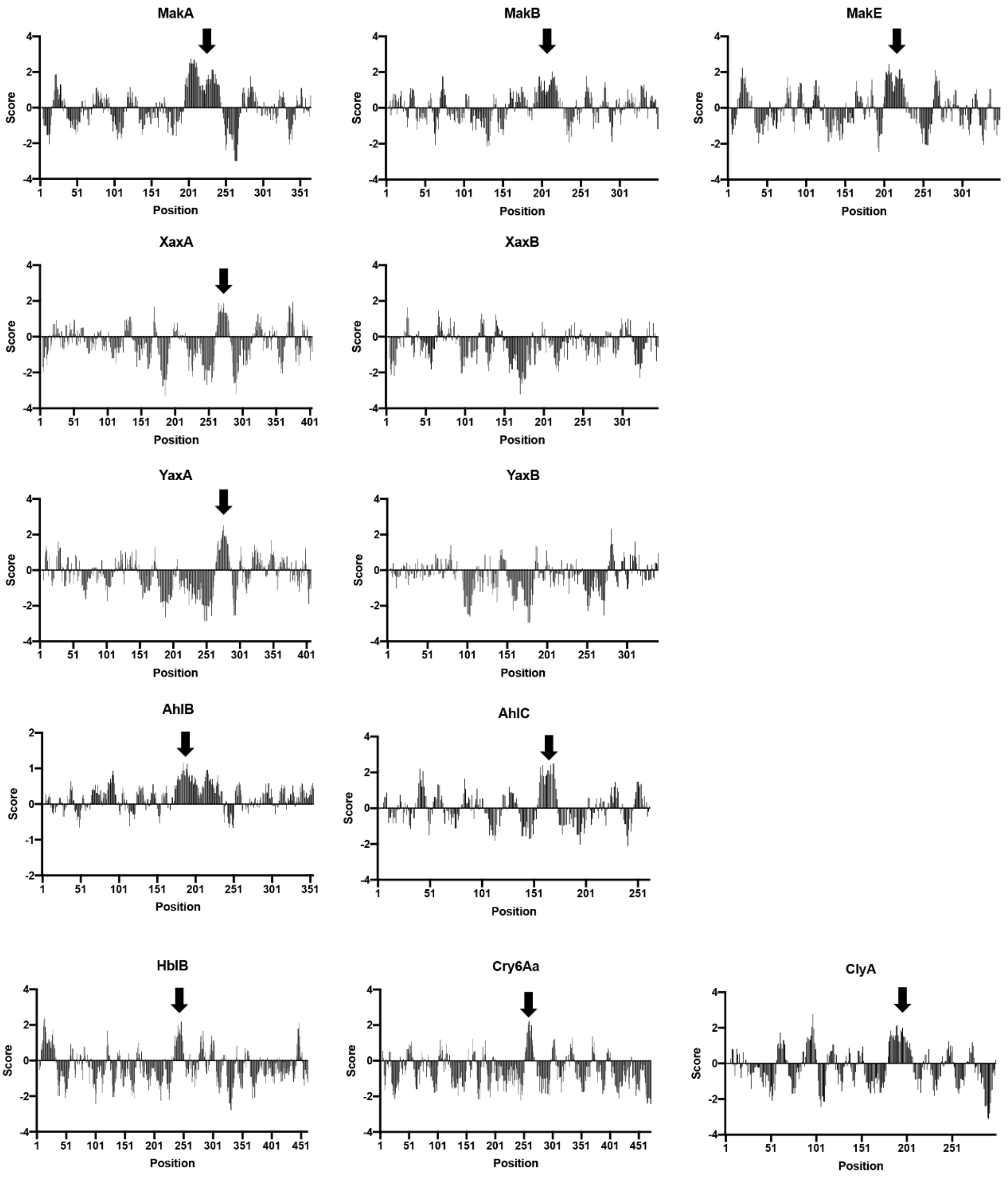

**Supplementary Fig. 6: Kyte and Doolittle hydropathy plots.**

Kyte and Doolittle plots were generated using ProtScale for structurally related proteins to MakA, MakB and MakE, identified by the Phyre2 database. The black arrow indicates predicted transmembrane regions.

#### Supplementary information

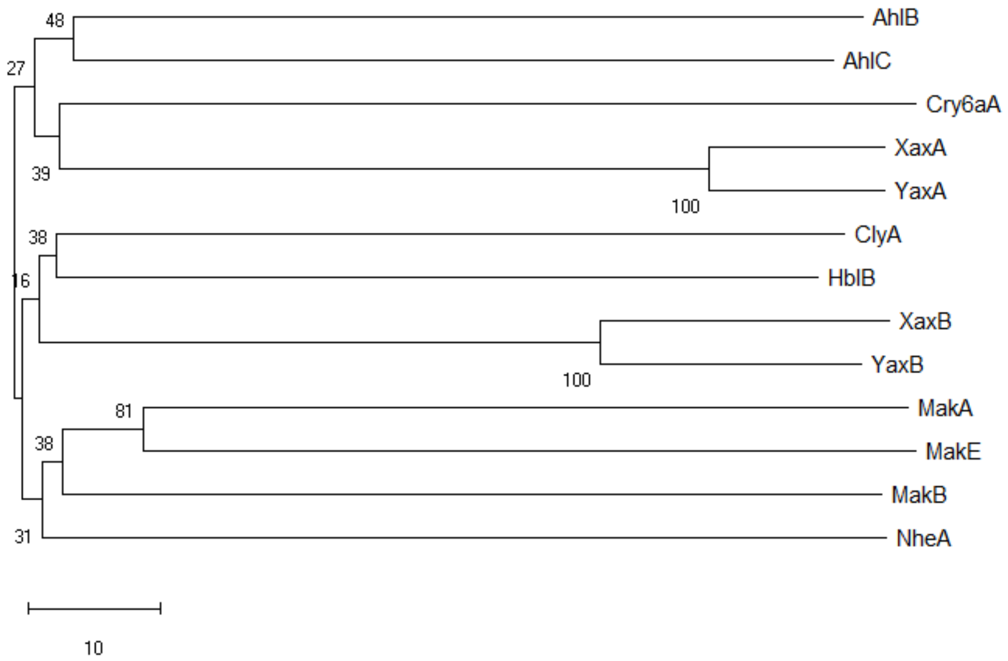

**Supplementary Fig. 7. Evolutionary relationships of Mak and other bacterial toxins.** Dendrogram showing hierarchical relationships between the structurally similar MakA, MakB, MakE and other ClyA family protein toxins using the Neighbor-Joining method (the bootstrap test was done with 500 replicates).

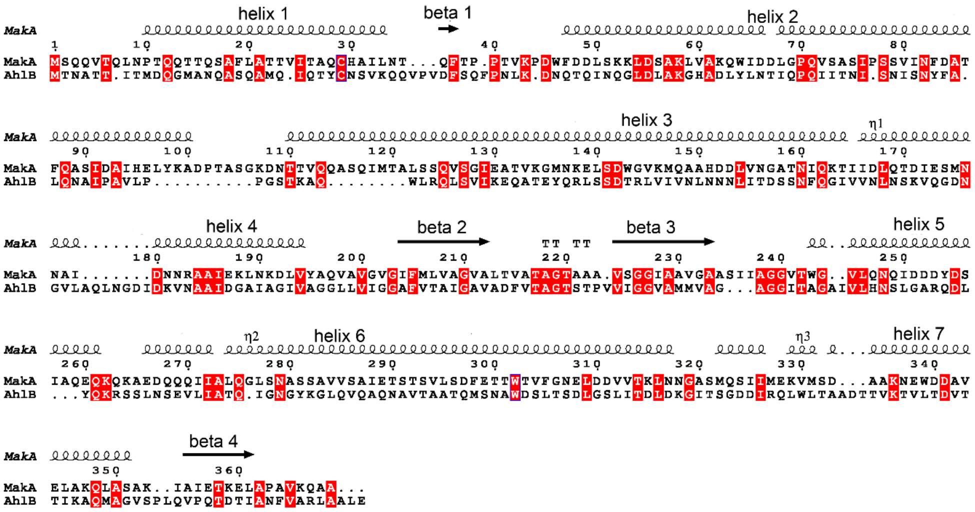

**Supplementary Fig. 8: Amino acids sequence alignment of MakA and AhlB.** Identical residues are highlighted in red. Secondary structure elements, based on the MakA structure, are indicated above the sequences.

#### Supplementary information

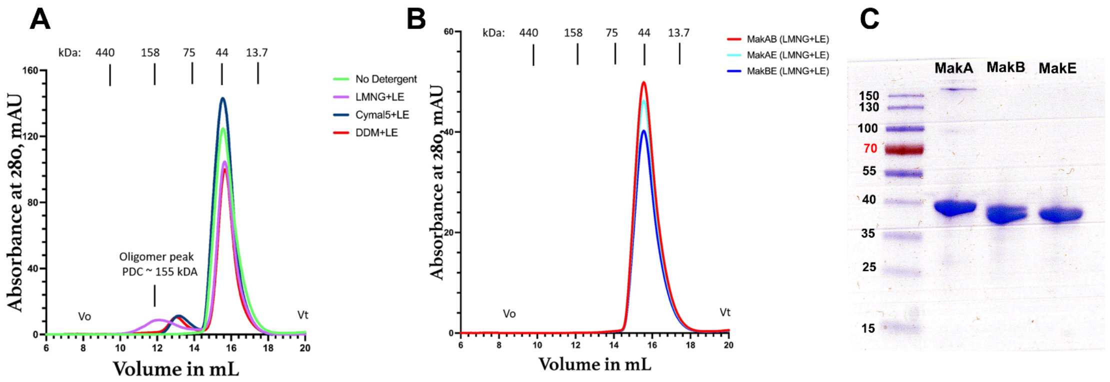

**Supplementary Fig. 9**: **Analysis of MakA/B/E proteins.**

***(A)*** Gel filtration analysis of an equimolar (250 nM) mixture of MakA/B/E proteins incubated in absence or in the presence of selected detergents. *Black* (no detergent), *Blue* (DDM+LE), *Grey* (cymal5+LE), *Green* (LMNG+LE). Upon addition of DDM+LE and cymal5+LE, the monomeric MakA/B/E proteins immediately associated, leading to a shift in the elution volume from 15.6 mL to 13.0 mL. The LMNG+LE detergent-bound MakA/B/E monomers slowly converted to higher order oligomers eluting at 12.1 mL. As shown in the corresponding gel filtration trace (*Green*), the LMNG+LE resulted in a predominant oligomer peak at approximately 155 kDa. $V_0$ indicates void volume- $V_t$ indicates the total liquid volume of the gel filtration column.

***(B)*** Gel filtration analysis of bipartite combinations (250 mM protein) performed at in buffer +LMNG+LE.

***(C)*** SDS PAGE and Coomassie Blue staining analysis of the recombinant MakA/B/E proteins used in this study.

#### Supplementary information

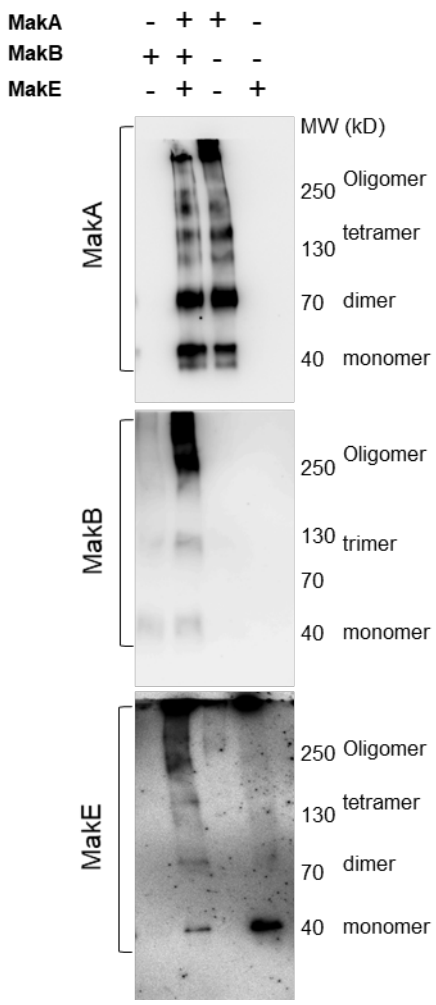

**Supplementary Fig. 10: The MakA/B/E tripartite oligomerizes after binding with liposomes**

Liposomes prepared from *E. coli* total lipid extract were incubated with MakA/BE tripartite (1:1:1, 500nM each protein) or its individual components, MakA, MakB, or MakE (500 nM). The pull-down assay including a crosslinking step was performed as described in Materials and Methods. The liposome bound proteins were detected with Mak protein-specific antiserum. All proteins are detected in oligomeric form when all proteins were incubated with liposomes together. MakA binds to the liposomes on its own and polymerizes. MakB does not interact with liposomes in the absence of the other Mak proteins. MakE interacts with the membrane by itself but does not form oligomers.

#### Supplementary information

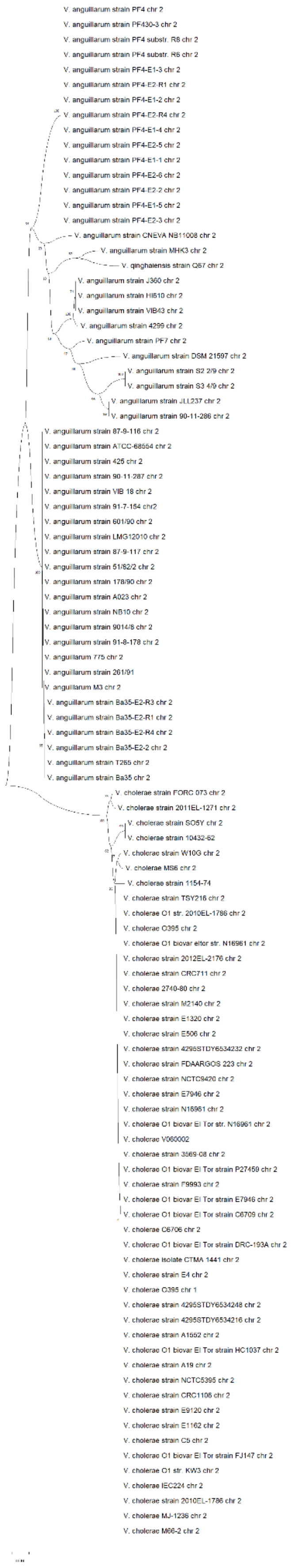

**Supplementary Fig. 11:** Phylogenetic tree based on the *mak* gene cluster sequences located on the genomes of the Vibrio species shown in **Supplementary information Table S2**. The tree was constructed using the Maximum Likelihood method. The number of each node represents the bootstrap value derived from 500 replicates.

### Supplementary information

#### Supplementary Table 1. Data collection and refinement statistics

| | MakE SeMet | MakE native | MakB SeMet | MakB native |
|---|---|---|---|---|
| **Data collection** | | | | |
| Space group | $P2_12_12_1$ | $P2_1$ | $P2_12_12_1$ | $P2_12_12_1$ |
| Cell dimensions | | | | |
| *a*, *b*, *c* (Å) | 66.6, 94.3, 116.6 | 56.9, 94.9, 63.6 | 53,2 54.2, 120.7 | 53,3 54.2, 120.8 |
| $\alpha, \beta, \gamma$ (°) | 90, 90, 90 | 90, 110.4, 90 | 90, 90, 90 | 90, 90, 90 |
| Resolution (Å) * | 49.3-2.60 (2.71-2.60) | 49.2-1.98 (2.05-1.98) | 49.5-1.92 (1.99-1.92) | 48.8-2.10 (2.16-2.10) |
| $R_{merge}$ | 0.141 (1.652) | 0.0.086 (0.998) | 0.075 (1.951) | 0.097 (0.860) |
| $I / \sigma I$ | 17.9 (2.0) | 9.6 (1.2) | 15.7 (1.4) | 9.0 (1.9) |
| Completeness (%) | 99.6(90.5) | 99.9(100) | 100(100) | 99.8(99.5) |
| Redundancy | 25.2(25.4) | 4.5(4.4) | 13.0(13.3) | 6.5(6.5) |
| CC1/2 | 1.00(0.836) | 1(0.572) | 0.999(0,778) | 0.995(0.816) |
| Molecules in a.u. | 2 | 2 | 1 | 1 |
| **Refinement** | | | | |
| Resolution (Å) | | 49.2-1.98 | | 60.41-2.10 |
| No. reflections (work/test) | | 44086(2000) | | 20016(1056) |
| $R_{work}$ / $R_{free}$ | | 0.177(0.235) | | 0.215(0.261) |
| No. atoms | | | | |
| Protein | | 5371 | | 5474 |
| Ligand/ion | | 14 ($4Ni^{2+}$, $2SO_4^{2-}$) | | 10 (2 $SO_4^{2-}$) |
| Water | | 412 | | 61 |
| *B*-factors (Å$^2$) | | | | |

#### Supplementary information

| | | | | |
|---|---|---|---|---|
| Protein | | 41.3 | | 48.4 |
| Ligand/ion | | 79.4 | | 61.04 |
| Water | | 43.1 | | 48.36 |
| R.m.s. deviations | | | | |
| Bond lengths (Å) | | 0.013 | | 0.003 |
| Bond angles (°) | | 1.14 | | 0.55 |
| PDB code | | 6TAO | | 6T8D |

For each structure, diffraction data were collected on a single crystal.

*Values in parentheses are for the highest-resolution shell.

#### Supplementary information

**Supplementary Table 2**. *Vibrio* strains with completely sequenced genomes that have the *mak* gene cluster

| [1]*Vibrio* Strain<br><br>O serogroup & biovar if known | [2]Chromosome | Coordinates in the genome | [3]Orientation of *mak* genes | [4]Accession number |
|---|---|---|---|---|
| *V. cholerae* O1 biovar El Tor strain A1552 | Chr. 2 | 832257-837904 | → | CP028895.1 |
| *V. cholerae* O1 biovar El Tor strain C6709 | Chr. 2 | 832241-837888 | → | CP047298.1 |
| *V. cholerae* O1 biovar El Tor strain P27459 | Chr. 2 | 835429-841076 | → | CP047300.1 |
| *V. cholerae* O1 biovar El Tor strain DRC-193A | Chr. 2 | 807261-812908 | → | CP047302.1 |
| *V. cholerae* O1 biovar El Tor strain E7946 | Chr. 2 | 832887-838534 | → | CP047304.1 |
| *V. cholerae* O1 biovar El Tor strain E7946 | Chr. 2 | 830081-835728 | → | CP024163.1 |
| *V. cholerae* O1 biovar El Tor strain CTMA_1441 | Chr. 2 | 821464-827198 | → | CP047060.1 |
| *V. cholerae* strain F9993 | Chr. 2 | 956732-962379 | ← | CP046841.1 |
| *V. cholerae* O1 biovar El Tor strain C6706 | Chr. 2 | 667286-672933 | ← | CP046845.1 |
| *V. cholerae* O1 biovar El Tor strain 3569-08 | Chr. 2 | 174549-180196 | ← | CP046743.1 |
| *V. cholerae* O1 biovar El Tor strain N16961 | Chr. 2 | 831898-837545 | → | LT906615.1 |
| *V. cholerae* O1 biovar El Tor strain N16961 | Chr. 2 | 831895-837542 | → | AE003853.1 |
| *V. cholerae* O1 biovar El Tor strain N16961 | Chr. 2 | 834209-839857 | → | CP028828.1 |
| *V. cholerae* O1 biovar El Tor strain E4 | Chr. 2 | 771425-777072 | ← | CP033513.1 |

| | | | | |
|---|---|---|---|---|
| *V. cholerae* O1 biovar El Tor strain V060002 | Chr. | 2149839-2155486 | ← | AP018677.1 |
| *V. cholerae* O139 biovar El Tor strain 4295STDY6534248 | Chr. 2 | 703431-709078 | → | LT992493.1 |
| *V. cholerae* O139 biovar El Tor strain 4295STDY6534232 | Chr. 2 | 1013978-1019625 | ← | LT992489.1 |
| *V. cholerae* O139 biovar El Tor strain 4295STDY6534216 | Chr. 2 | 481479-487126 | ← | LT992487.1 |
| *V. cholerae* O1 biovar El Tor strain FDAARGOS_223 | Chr. 2 | 347303-352950 | ← | CP020407.2 |
| *V. cholerae* O1 biovar El Tor strain HC1037 | Chr. 2 | 532359-538006 | → | CP026648.1 |
| *V. cholerae* O1 biovar El Tor strain A19 | Chr. 2 | 831898-837545 | → | LT907990.1 |
| *V. cholerae* O1 biovar El Tor strain NCTC9420 | Chr. 2 | 827597-833244 | → | CP013320.1 |
| *V. cholerae* O1 biovar El Tor strain NCTC5395 | Chr. 2 | 892952-898599 | → | CP013318.1 |
| *V. cholerae* O1 biovar El Tor strain M2140 | Chr. 2 | 618070-623717 | → | CP013316.1 |
| *V. cholerae* O1 biovar El Tor strain E9120 | Chr. 2 | 864606-870253 | → | CP013314.1 |
| *V. cholerae* O1 biovar El Tor strain E1320 | Chr. 2 | 108643-114290 | → | CP013312.1 |
| *V. cholerae* O1 biovar El Tor strain E1162 | Chr. 2 | 992794-998441 | → | CP013310.1 |
| *V. cholerae* O1 biovar El Tor strain E506 | Chr. 2 | 852260-857907 | → | CP013308.1 |
| *V. cholerae* O1 biovar El Tor strain CRC1106 | Chr. 2 | 859687-865334 | → | CP013306.1 |
| *V. cholerae* O1 biovar El Tor strain CRC711 | Chr. 2 | 823844-829491 | → | CP013304.1 |

#### Supplementary information

| | | | | |
|---|---|---|---|---|
| *V. cholerae* O1 biovar El Tor strain C5 | Chr. 2 | 865423-871070 | → | CP013302.1 |
| *V. cholerae* O1 biovar El Tor strain 2740-80 | Chr. 2 | 861068-866715 | → | CP016325.1 |
| *V. cholerae* O1 strain KW3 | Chr. 2 | 807251-812898 | → | CP006948.1 |
| *V. cholerae* O1 biovar El Tor strain TSY216 | Chr. 2 | 809852-815499 | → | CP007654.1 |
| *V. cholerae* O1 biovar El Tor strain FJ147 | Chr. 2 | 806032-811679 | → | CP009041.1 |
| *V. cholerae* O1 biovar El Tor strain 2012EL-2176 | Chr. 2 | 533667-539314 | → | CP007635.1 |
| *V. cholerae* O1 strain IEC224 | Chr. 2 | 831720-837367 | → | CP003331.1 |
| *V. cholerae* O1 biovar El Tor strain 2010EL-1786 | Chr. 2 | 532427-538074 | → | CP003070.1 |
| *V. cholerae* O1 biovar El Tor strain MJ-1236 | Chr. 2 | 508290-513937 | ← | CP001486.1 |
| *V. cholerae* O1 classical strain O395 | Chr. 2 | 870719-876366 | → | CP001236.1 |
| *V. cholerae* O1 classical strain O395 | Chr. 2 | 387576-393223 | ← | CP000626.1 |
| *V. cholerae* O1 classical strain O395 | Chr. 1 | 387576-393310 | ← | CP045718.1 |
| *V. cholerae* O1 biovar El Tor strain M66-2 | Chr. 2 | 805881-811528 | → | CP001234.1 |
| *V. cholerae* non-O1/O139 environmental isolate strain W10G | Chr. 2 | 788603-794251 | → | CP053795.1 |
| *V. cholerae* O1 biovar El Tor strain 2010EL-1786 | Chr. 2 | 664915-670552 | → | CP038179.2 |
| *V. cholerae* O1 biovar El Tor strain MS6 | Chr. 2 | 847066-852712 | → | AP014525.1 |

| | | | | |
|---|---|---|---|---|
| *V. cholerae* non-O1/O139 environmental isolate strain SO5Y | Chr. 2 | 814545-820160 | → | CP053801.1 |
| *V. cholerae* O27 strain 10432-62 | Chr. | 338019-343642 | ← | CP010812.1 |
| *V. cholerae* strain FORC_073 | Chr. 2 | 759334-765000 | → | CP024083.1 |
| *V. cholerae* O1 strain 2011EL-1271 | Chr. 2 | 386300-391301 | ← | CP046838.1 |
| *V. cholerae* O49 strain 1154-74 | Chr. | 2336425-2341366 | ← | CP010811.1 |
| *V. anguillarum* O1 strain 425 | Chr. 2 | 790251-794855 | ← | CP020533.1 |
| *V. anguillarum* O1 strain 87-9-116 | Chr. 2 | 7345-11949 | ← | CP021981.1 |
| *V. anguillarum* O1 strain VIB 18 | Chr. 2 | 543776-548380 | ← | CP011437.1 |
| *V. anguillarum* O1 strain 90-11-287 | Chr. 2 | 540749-545353 | ← | CP011476.1 |
| *V. anguillarum* O1 strain 91-7-154 | Chr. 2 | 540911-545515 | ← | CP010083.1 |
| *V. anguillarum* O1 strain 601/90 | Chr. 2 | 540787-545391 | ← | CP010077.1 |
| *V. anguillarum* O1 strain 178/90 | Chr. 2 | 540837-545441 | ← | CP011471.1 |
| *V. anguillarum* O1 strain LMG12010 | Chr. 2 | 536719-541323 | ← | CP011469.1 |
| *V. anguillarum* O1 strain 87-9-117 | Chr. 2 | 538384-542988 | ← | CP010047.1 |
| *V. anguillarum* O1 strain 51/82/2 | Chr. 2 | 539273-543877 | ← | CP010043.1 |
| *V. anguillarum* O1 strain 9014/8 | Chr. 2 | 539619-544223 | ← | CP010039.1 |
| *V. anguillarum* O1 strain A023 | Chr. 2 | 541898-546502 | ← | CP010037.1 |

| | | | | |
|---|---|---|---|---|
| *V. anguillarum* O1 strain 91-8-178 | Chr. 2 | 543326-547930 | ← | CP010035.1 |
| *V. anguillarum* O1 strain NB10 | Chr. 2 | 7344-11948 | ← | LK021129.1 |
| *V. anguillarum* O1 strain M3 | Chr. 2 | 543797-548401 | ← | CP006700.1 |
| *V. anguillarum* O1 strain 775 | Chr. 2 | 543821-548425 | ← | CP002285.1 |
| *V. anguillarum* O1 strain Ba35-E2-R3 | Chr. 2 | 542397-547001 | ← | CP031528.1 |
| *V. anguillarum* O1 strain Ba35-E2-R1 | Chr. 2 | 544119-548723 | ← | CP031524.1 |
| *V. anguillarum* O1 strain Ba35-E2-R4 | Chr. 2 | 542233-546837 | ← | CP031532.1 |
| *V. anguillarum* O1 strain Ba35-E2-2 | Chr. 2 | 543973-548577 | ← | CP031520.1 |
| *V. anguillarum* O1 strain T265 | Chr. 2 | 543438-548042 | ← | CP010041.1 |
| *V. anguillarum* O1 strain Ba35 | Chr. 2 | 543336-547940 | ← | CP010031.1 |
| *V. anguillarum* O3 strain PF4-E2-3 | Chr. 2 | 859280-863881 | → | CP031493.1 |
| *V. anguillarum* O3 strain PF4-E1-5 | Chr. 2 | 876563-881164 | → | CP031486.1 |
| *V. anguillarum* O3 strain PF4-E2-2 | Chr. 2 | 859282-863883 | → | CP031491.1 |
| *V. anguillarum* O3 strain PF4-E2-5 | Chr. 2 | 859284-863885 | → | CP031497.1 |
| *V. anguillarum* O3 strain PF4-E2-6 | Chr. 2 | 859284-863885 | → | CP031499.1 |
| *V. anguillarum* O3 strain PF4-E1-1 | Chr. 2 | 874235-878836 | → | CP031478.1 |
| *V. anguillarum* O3 strain PF4-E1-4 | Chr. 2 | 874234-878835 | → | CP031484.1 |

| | | | | |
|---|---|---|---|---|
| *V. anguillarum* O3 strain PF4-E2-R4 | Chr. 2 | 859284-863885 | → | CP031495.1 |
| *V. anguillarum* O3 strain PF4-E1-2 | Chr. 2 | 874235-878836 | → | CP031480.1 |
| *V. anguillarum* O3 strain PF4-E2-R1 | Chr. 2 | 859287-863888 | → | CP031489.1 |
| *V. anguillarum* O3 strain PF4-E1-3 | Chr. 2 | 874234-878835 | → | CP031482.1 |
| *V. anguillarum* O3 strain PF4 substr. R6 | Chr. 2 | 860025-864626 | → | CP023432.1 |
| *V. anguillarum* O3 strain PF4 substr. R8 | Chr. 2 | 860021-864622 | → | CP023292.1 |
| *V. anguillarum* O3 strain PF4 | Chr. 2 | 860022-864623 | → | CP023290.1 |
| *V. anguillarum* O3 strain PF4 substr. R4 | Chr. 2 | 860025-864626 | → | CP023288.1 |
| *V. anguillarum* O3 strain PF430-3 | Chr. 2 | 577775-582379 | ← | CP011467.1 |
| *V. anguillarum* O3 strain PF7 | Chr. 2 | 343423-348025 | → | CP011465.1 |
| *V. anguillarum* O2 strain J360 | Chr. 2 | 175977-180582 | ← | CP034673.1 |
| *V. anguillarum* O1 strain VIB43 | Chr. 2 | 7356-11961 | ← | CP023055.1 |
| *V. anguillarum* O2a strain HI610 | Chr. 2 | 582185-586790 | ← | CP011463.1 |
| *V. anguillarum* non-O1/O2/O3/O4/O5 strain MHK3 | Chr. 2 | 324177-328777 | → | CP022469.1 |
| *V. anguillarum* O3 strain CNEVA NB11008 | Chr. 2 | 7355-11962 | ← | CP022104.1 |
| *V. anguillarum* O2b strain 4299 | Chr. 2 | 542862-547467 | ← | CP011459.1 |
| *V. anguillarum* O1 strain S2 2/9 | Chr. 2 | 529358-533980 | ← | CP011473.1 |

### Supplementary information

| | | | | |
|---|---|---|---|---|
| *V. anguillarum* O1 strain S3 4/9 | Chr. 2 | 7355-11977 | ← | CP022100.1 |
| *V. anguillarum* O2 strain DSM 21597 | Chr. 2 | 576229-580860 | ← | CP010085.1 |
| *V. anguillarum* O1 strain JLL237 | Chr. 2 | 7354-11220 | ← | CP022102.1 |
| *V. anguillarum* O1 strain 90-11-286 | Chr. 2 | 714386-718252 | ← | CP011461.1 |
| *V. anguillarum* O1 strain 261/91 | Chr. 2 | 544444-547945 | ← | CP010033.1 |
| *V. anguillarum* O1 strain ATCC-68554 | Chr. 2 | 983079-989192 | → | CP023209.1 |
| *V. qinghaiensis* strain Q67 | Chr. 2 | 28543-33164 | ← | CP022742.1 |

1. Strains represented more than once in the table have undergone repeated sequencing of their genomes and were reported independently. Identification of the *mak* gene cluster was therefore indicated for each report using the corresponding accession number.
2. Abbreviations: Chr. 1 = chromosome 1; Chr. 2 = chromosome 2; Chr. = genome composed of only one chromosome.
3. → indicates that the gene cluster is oriented in the clockwise direction relative to the replication origin and ← indicates that the gene cluster is oriented in the counterclockwise direction.
4. GenBank; https://www.ncbi.nlm.nih.gov/genbank/

### Supplementary information

##### Supplementary Table 3: Bacterial strains and plasmids used in this study

| Strain/ plasmid | Description/relevant characteristics | Reference / Source |
|---|---|---|
| ***Vibrio cholerae*** | | |
| ***Vibrio cholerae*** A1552: serogroup O1, biotype El Tor, Rif$^r$ | | Yildiz et al., 1998(26) |
| MDS002 | A1552Δ*makD* | Dongre et al., 2018(1) |
| MDS003 | A1552Δ*makC* | Dongre et al., 2018(1) |
| MDS004 | A1552Δ*makB* | Dongre et al., 2018(1) |
| MDS005 | A1552Δ*makA* | Dongre et al., 2018(1) |
| MDC008 | A1552Δ*makE* | This study |
| MDS013 | A1552Δ*flhA* | Dongre et al., 2018(1) |
| TAKA-23 | A1552Δ*flgT* | This Study |
| MDC003 | A1552Δ*makB*/p*makB* | Dongre et al., 2018(1) |
| MDC002 | A1552Δ*makC*/p*makC* | Dongre et al., 2018(1) |
| ***Escherichia coli*** | | |
| DH5α | *F$^-$ Φ80lacZΔM15 Δ(lacZYA-argF) U169 recA1 endA1 hsdR17($r_K^-$, $m_K^+$) phoA supE44 thi-1 gyrA96 relA1 λ$^-$* | Grant et al., 1985(27) |
| BL21(DE3) | *fhuA*2 [*lon*] *ompT gal* (λ DE3) [dcm] Δ*hsdS*<br><br>λ DE3 = λ s*BamHI*o Δ*EcoRI-B* int::(*lacI*::PlacUV5::*T7 gene1*) i21Δ*nin5* | Studier and Moffatt,1986(28) |
| SM10λpir | *thi thr leu tonA lacY supE recA*::RP4-2 Tc::Mu Km λpir | Miller et al. 1987(29) |
| MDS611 | Top10/pBAD18; Ap$^r$ | Dongre et al., 2018(1) |
| AA1 | Top10/p*makB$^+$A$^+$E$^+$* | This Study |
| AA2 | Top10/p*makB$^+$A$_{L193T,L246T}$E$^+$* | This Study |
| AA3 | Top10/p*makB$^+$A$_{L193T,L207T,L213T,L246T}$E$^+$* | This Study |
| **Plasmids** | | |
| pCVD442 | Suicide plasmid; R6K ori, mobRP4, bla, *sacB*; Ap$^r$ | Donnenberg et al. 1991(3) |

### Supplementary information

| | | |
|---|---|---|
| pMDC008 | pCVD442::Δ*makE* | This Study |
| pTAKA-23 | pCVD442::Δ*flgT* | This Study |
| pMDC003 | pBAD18-*makB* | Dongre et al., 2018(1) |
| pMDC002 | pBAD18-*makC* | Dongre et al., 2018(1) |
| pAA1 | pBAD18- $makB^{+}A^{+}E^{+}$ | This Study |
| pAA2 | pBAD18-$makB^{+}A_{L193T,L246T}E^{+}$ | This Study |
| pAA3 | pBAD18-$makB^{+}A_{L193T,L207T,L213T,L246T}E^{+}$ | This Study |

### Supplementary information

#### Supplementary Table 4. Primers used in this study

| Sequence 5'→3' | Restriction site | For construction of |
|---|---|---|
| atagagctccctcgttatttcactgtagag | makB-pBAD18-5Sac1 | *makB*$^+$*A*$^+$*E*$^+$ |
| gtctctagattacttgatttgagcttttgttat | makE-pBAD18-3Xba1 | *makB*$^+$*A*$^+$*E*$^+$ |
| gtagtcatcatcaatttgattttgcgttacgccccatgt aacgccaccag | makA_t736a_t737c_a738g_F | *makB*$^+$*A*$_{L193T,L246T}$*E*$^+$ |
| ctggtggcgttacatggggcgtaacgcaaaatca aattgatgatgactac | makA_t736a_t737c_a738g_R | *makB*$^+$*A*$_{L193T,L246T}$*E*$^+$ |
| caacgcctgcgacggtcataaagatgccaacac caac | makA_c619a_t620c_F | *makB*$^+$*A*$_{L193T,L246T}$*E*$^+$ |
| gttggtgttggcatctttatgaccgtcgcaggcgttg | makA_c619a_t620c_R | *makB*$^+$*A*$_{L193T,L246T}$*E*$^+$ |
| gtagtcatcatcaatttgattttgcgttacgccccatgt aacgccaccag | makA_t736a_t737c_a738g_F | *makB*+*A*$_{L193T,L207T,L213T,L246T}$*E*+ |
| ctggtggcgttacatggggcgtaacgcaaaatca aattgatgatgactac | makA_t736a_t737c_a738g_R | *makB*+*A*$_{L193T,L207T,L213T,L246T}$*E*+ |
| ccgcaacttgtgcatagaccgtatctttattcagtttct caatgg | makA_c577a_t578c_F | *makB*+*A*$_{L193T,L207T,L213T,L246T}$*E*+ |
| ccattgagaaactgaataaagatacggtctatgca caagttgcgg | makA_c577a_t578c_R | *makB*+*A*$_{L193T,L207T,L213T,L246T}$*E*+ |
| cggttgcgacggtagtcgcaacgcctgcga | makA_c637a_t638c_F | *makB*+*A*$_{L193T,L207T,L213T,L246T}$*E*+ |
| tcgcaggcgttgcgactaccgtcgcaaccg | makA_c637a_t638c_R | *makB*+*A*$_{L193T,L207T,L213T,L246T}$*E*+ |
| caacgcctgcgacggtcataaagatgccaacac caac | makA_c619a_t620c_F | *makB*+*A*$_{L193T,L207T,L213T,L246T}$*E*+ |
| gttggtgttggcatctttatgaccgtcgcaggcgttg | makA_c619a_t620c_R | *makB*+*A*$_{L193T,L207T,L213T,L246T}$*E*+ |
| cattctagaagctgcagtctccggcggtatc | makE-A F | *ΔmakE* |
| cccatccactataaactaacaagaatagaaggg ggcgtaacc | makE-B R | *ΔmakE* |
| tgttagtttatagtggatgggaagtaattgatttcgct ccag | makE-C F | *ΔmakE* |
| cattctagactgatgggaaagctcattaaac | makE-D R | *ΔmakE* |
| cgctctagatgctgcgtcacttggattgaa | flgT-A F | *ΔflgT* |

| | | |
|---|---|---|
| cccatccactataaactaacatgcatgagccgatg gaagttgcaatctatgcagcggctaaga | flgT-B R | *ΔflgT* |
| tgttagtttatagtggatgggattcaaatcggcgatg tgatg | flgT-C F | *ΔflgT* |
| cgctctagaacagatattccgattgacctgcaggtc aatcggaatatctgt | flgT-D R | *ΔflgT* |
| gtaacggtctcccatgttaagcccaaatgttgatac | MakB-F | Cloning of recombinant MakB |
| gttacctcgagttactgttcttgaatggtgttatttag | MakB-R | Cloning of recombinant MakE |
| gtaactcatgaatcaatcagccagcgag | MakE-F | Cloning of recombinant MakE |
| gtaacaagcttacttgatttgagcttttgttatttg | MakE-R | Cloning of recombinant MakE |

### Supplementary information
